# Partial redundancy buffers deleterious effects of mutating *DNA methyltransferase 1-1* (*MET1-1*) in polyploid wheat

**DOI:** 10.1101/2024.07.26.605257

**Authors:** Samuel Burrows, Delfi Dorussen, Joseph Crudgington, Giorgia Di Santolo, James Simmonds, Marco Catoni, Philippa Borrill

## Abstract

DNA methylation is conserved across biological kingdoms, playing important roles in gene expression, transposable element silencing and genome stability. Altering DNA methylation could generate additional phenotypic variation for crop breeding, however the lethality of epigenetic mutants in crop species has hindered its investigation. Here, we exploit partial redundancy between homoeologs in polyploid wheat to generate viable mutants in the DNA *methyltransferase 1-1* (*MET1-1*) gene with altered methylation profiles. In both *Triticum turgidum* (tetraploid wheat) and *Triticum aestivum* (hexaploid wheat) we identified clear segregation distortions of higher-order mutants (5/6 and 6/6 mutant *met1-1* copies in hexaploid and 3/4 and 4/4 copies in tetraploid) when genotyping segregating seeds and seedlings, which we attribute to reduced transmission of null mutant gametes. We found that the reduced transmission occurred from both the maternal and paternal gametes, however, we did not detect any deleterious effects on pollen development. The loss of four or more functional copies of *MET1-1* results in decreased CG methylation in hexaploid wheat. Changes to gene expression increase stepwise with the number of mutant alleles suggesting a dosage dependent effect. Finally, we identify heritable changes to flowering and awn phenotypes which segregate independently of *MET1-1*. Together our results demonstrate that polyploidy can be leveraged to generate quantitative changes to CG methylation without the lethal consequences observed in other crops, opening the potential to exploit novel epialleles in plant breeding.

## Introduction

DNA methylation on cytosine bases is an epigenetic modification which is conserved across biological kingdoms and plays important roles in gene expression, transposable element silencing and genome stability. In plants, cytosine methylation can occur in any context (CG, CHG and CHH, where H is A, T, or C) although CG methylation is the most prevalent context in plant species examined to date (Feng *et al*., 2010; Niederhuth *et al*., 2016; Domb *et al*., 2020; Li *et al*., 2023). Methylation is established and maintained by multiple enzymes which function in specific or overlapping contexts. CG methylation is maintained during cell division by the methyltransferase 1 (MET1) which recognises hemi-methylated CG sites following DNA replication and methylates the opposite strand (Kankel *et al*., 2003; Zhang *et al*., 2018).

Disruption of the *MET1* gene in Arabidopsis (*Arabidopsis thaliana*), tomato (*Solanum lycopersicum*) and rice (*Oryza sativa*) has revealed major differences in the phenotypic impact and severity of null mutations between species. Arabidopsis *met1* mutants, including complete and partial loss-of-function alleles, exhibit dwarfism, delayed flowering, abnormal embryo development and seed abortion (Kankel *et al*., 2003; Saze *et al*., 2003; Xiao *et al*., 2006; Srikant *et al*., 2022). Null *met1* alleles have severe consequences with segregation distortion resulting in an underrepresentation of homozygous mutants, which are only occasionally viable to maturity (Saze *et al*., 2003; Srikant *et al*., 2022). In rice, null homozygous mutants were formed at the expected ratio but underwent necrotic death at seedling stage (Hu *et al*., 2014) and in tomato, homozygous null mutants could not be recovered (Yang *et al*., 2019). These species also vary in CG methylation loss in *met1* mutants, with a stronger reduction in Arabidopsis than in rice (98.3% vs 75.7%) and only a 25% reduction in tomato CRISPR mutants, although this may be due to chimerism (Cokus *et al*., 2008; Hu *et al*., 2014; Yang *et al*., 2019).

Differences in the segregation and viability of homozygous null mutants may be due to variation in epigenetic backup systems: in Arabidopsis these backup systems operate through changes to the RNA-directed DNA methylation (RdDM) process, changes to expression of DNA demethylases and progressive H3K9 remethylation of heterochromatin (Mathieu *et al*., 2007). Stochasticity in these processes explains the rarity of homozygous null mutants (Mathieu *et al*., 2007). In rice, a duplicate copy of *MET1* has been proposed to provide a back-up mechanism for essential CG methylation, although the lethality of homozygous mutants at seedling stage indicates incomplete compensation (Hu *et al*., 2014; Yamauchi *et al*., 2014).

Mutants in *met1* have not been examined within a polyploid species where duplicate gene copies could provide a back-up mechanism. Alternatively, gene copies could be dosage-sensitive with a mutation in just one copy of *met1* affecting DNA methylation. The gene balance hypothesis proposes that dosage-sensitive genes are generally involved in macromolecular complexes or are regulatory genes such as transcription factors, rather than enzymes like *MET1* (Veitia & Birchler, 2010; Birchler & Veitia, 2012). Moreover, it is generally expected that mutations in enzymes would be recessive, making it unlikely to find phenotypic effects in a partial *met1* mutant in a polyploid (Kondrashov & Koonin, 2004).

However, some enzymes are dosage-sensitive in polyploids, such as the E3 ubiquitin ligase gene *GW2* which has a dosage effect on grain width in wheat (Wang *et al*., 2018). MET1 has been reported to interact at the protein level with histone deacetylase HDA6 and MEDEA in Arabidopsis (Liu *et al*., 2012; Schmidt *et al*., 2013), making a dosage effect more likely since it could be part of a macromolecular complex. Opposing this view, heterozygous mutants in tomato and rice maintained similar CG methylation to wild type plants (Hu *et al*., 2014; Yang *et al*., 2019), indicating only one allele was sufficient to compensate, and the same was found for some Arabidopsis accessions, although CG methylation was reduced up to 64% in others (Srikant *et al*., 2022). Therefore, it is difficult to predict whether a partial mutant in *MET1* in a polyploid species would induce changes to CG methylation.

If *MET1* is dosage-sensitive in polyploids, partial *met1* mutants could be used to alter methylation status and produce stable epialleles for use in plant breeding. Epialleles affecting diverse traits have been identified in a range of plant species, including alterations in floral morphology, fruit ripening, plant height and climate adaptation (Cubas *et al*., 1999; Manning *et al*., 2006; Miura *et al*., 2009; He *et al*., 2018; Srikant & Tri Wibowo, 2021). The deliberate induction of stable epialleles has been demonstrated through epiRIL populations generated genetically using *met1* and *ddm1* (*Decreased DNA Methylation 1*) mutants in Arabidopsis (Johannes *et al*., 2009; Reinders *et al*., 2009; Zhang *et al*., 2013; Catoni & Cortijo, 2018). The potential to exploit novel epialleles in plant breeding has been proposed, but the production of epiRIL populations in crop species as a foundation to generate novel epialleles has been limited by the lethality of mutants (Springer & Schmitz, 2017). For example, no null mutants were identified in *MET1* orthologs in crop species with large genomes including in a barley TILLING population (genome size of ∼5Gb) (Schreiber *et al*., 2019) and in maize TILLING and transposon populations (genome size of ∼2.4 Gb) (Li *et al*., 2014). This may be due the large numbers of transposable elements in large genomes causing strong genetic and developmental defects in *met1* mutants through their remobilisation (Mirouze & Vitte, 2014).

Here we explore whether it is possible to use a polyploid crop species to generate quantitative changes to CG methylation by knocking out differing numbers of *met1* alleles, without causing lethal consequences. We use polyploid wheat as a model system due to the availability of sequenced mutant populations which can be readily applied to breeding worldwide without legislative complications surrounding gene editing (Krasileva *et al*., 2017). We generate *met1* mutants in tetraploid (*Triticum turgidum*) and hexaploid (*Triticum aestivum*) wheat with up to three null alleles (of four) in tetraploid wheat, and five null alleles (of six) in hexaploid wheat. Contrary to the predictions of the gene dosage balance hypothesis (Veitia & Birchler, 2010) we find changes to global DNA methylation, transcriptional responses and effects on fertility and plant growth linked to mutant allele copy number. We also identify stably inherited phenotypic alterations. Our findings provide insights into using polyploidy to generate quantitative changes to CG methylation for epigenetic breeding in crops.

## Results

### Null *met1-1* mutants cannot be recovered in polyploid wheat

*METHYLTRANSFERASE 1* (*MET1*) genes were previously identified in hexaploid wheat (*Triticum aestivum*) on the group 2, 5, and 7 chromosomes, with the group 5 and 7 *MET1* homologs predicted to be pseudogenes (Figure 1A) (Thomas *et al*., 2014). The group 2 genes are orthologous to *MET1b* in rice, also known as *OsMET1-2,* which strongly affects CG methylation (Figure 1A) (Hu *et al*., 2014). The group 2 genes (*TraesCS2A02G235900*, *TraesCS2B02G260800*, *TraesCS2D02G241800*) are expressed in numerous above-ground tissues (Figure 1B), whereas expression of the group 5 and group 7 genes was virtually undetectable in most of these tissues (Supplementary Figure 1). We therefore concentrated our analysis on the group 2 genes which will be referred to as *MET1-A1*, *MET1-B1*, and *MET1-D1* in accordance with the gene nomenclature guidelines for wheat (Boden *et al*., 2023) and a previous study by Li *et al*. (2021).

**Figure 1.**
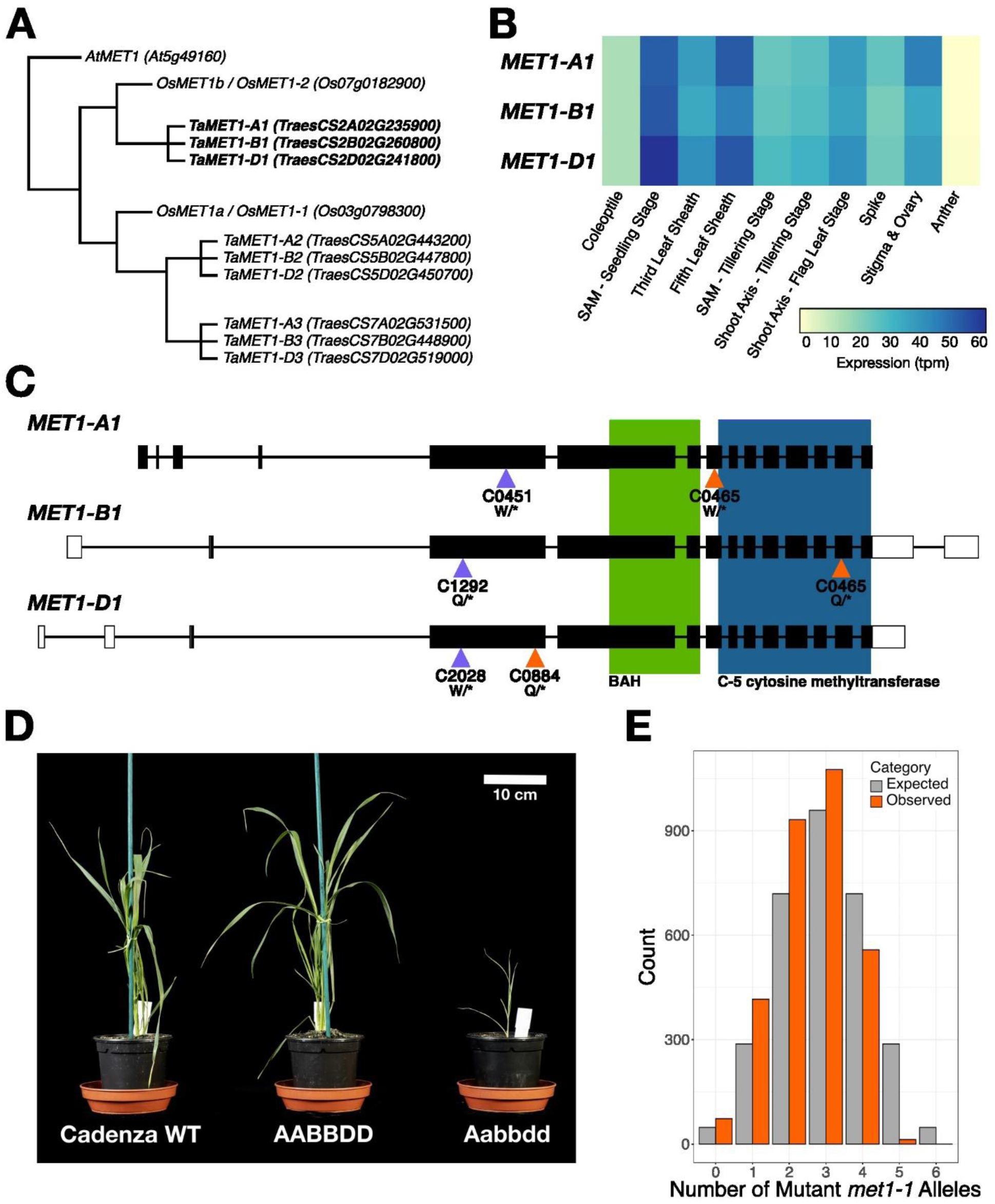
Null *met1-1* mutants cannot be produced by crossing TILLING lines with disruptions in the three *MET1-1* homoeologs. A) Evolutionary relationships between *MET1* homologs in Arabidopsis (*AtMET1*), rice (*OsMET1a* and *OsMET1b*), and wheat (homoeolog groups *TaMET1-1*, *TaMET1-2*, and *TaMET1-3*). B) Expression of the three *MET1-1* homoeologs (*TraesCS2A02G235900*, *TraesCS2B02G260800*, *TraesCS2D02G241800*) in above-ground tissues across development in the Chinese Spring cultivar. SAM = shoot apical meristem. C) Structure of the three *MET1-1* homoeologs. Filled rectangles represent exons, empty rectangles represent untranslated regions, and lines represent introns. Triangles indicate the positions of the premature termination codon mutations in the TILLING lines used to produce the two hexaploid F_2_ populations (C0465xC0884, orange, and C0451xC2028xC1292, purple). Green box signifies Bromo adjacent homology domain (BAH) and blue box signifies C-5 cytosine methyltransferase domain. D) Cadenza WT, WT segregant (AABBDD), and Aabbdd individuals from the C0465xC0884 segregating F_2_ population. E) Expected (grey) and observed (orange) number of individuals with each possible number of mutant *met1-1* alleles in C0465xC0884 segregating F_2_ population.

We produced independent segregating F_2_ populations in hexaploid wheat from which, in theory, all combinations of loss-of-function *met1-1* mutants could be recovered (Figure 1C, Supplementary Figure 2). However, we were unable to recover null *met1-1* mutants with homozygous loss-of-function mutations in all three homoeologs from either population, despite screening 3,068 individuals from the C0465xC0884 population, and 328 individuals from the C0451xC2028xC1292 population. Moreover, we only found one plant with five non-functional *MET1-1* alleles which was capable of germination, although its growth was severely stunted and the plant died before progressing to tillering (Zadoks growth stage 20) (Figure 1D). The genotype ratios observed in both populations were significantly different from expected Mendelian ratios as individuals with four or more non-functional *MET1-1* alleles were underrepresented (C0465xC0884 Figure 1E, χ^2^ p=1.63x10^-103^; C0451xC2028xC2192 Supplementary Table 1, χ^2^ p=5.05x10^-12^). We concluded that higher- order *met1-1* mutants have reduced viability in hexaploid wheat.

### Reduced viability of null *met1-1* gametes hinders recovery of higher-order *met1-1* mutants

We hypothesized that the absence of functional *MET1-1* alleles within a gamete might cause this segregation distortion. We developed a grain genotyping method using the endosperm- containing half of the grain (Figure 2A) which contains maternal to paternal DNA at a 2:1 ratio (endosperm originates from two female central cell nuclei being fertilised by one male sperm cell). This allowed us to determine which parent donated mutant alleles to the offspring (grain) by separating the heterozygous genotypes into two clusters (Figure 2B). We confirmed the accuracy of this method using reciprocal crosses for an independent mutation in the semi-dwarfing gene *Rht13* (Borrill *et al*., 2022) (Figure 2B). Grain genotypes were consistent with seedlings grown from the embryo-containing halves of the grain (Supplementary Table 2). We genotyped 1,710 grains from a segregating F_2_ hexaploid wheat population (C0465xC0884) and observed far fewer grains which inherited three mutant *met1- 1* alleles from one parent than expected (generalised linear model with log link, p<2x10^-16^; Figure 2C). We detected fewer grains which inherited three mutant alleles from the maternal side than from the paternal side (18 vs 32 grains). There was no reduction in the number of grains which inherited one or two mutant alleles from the same parent (Figure 2C).

**Figure 2.**
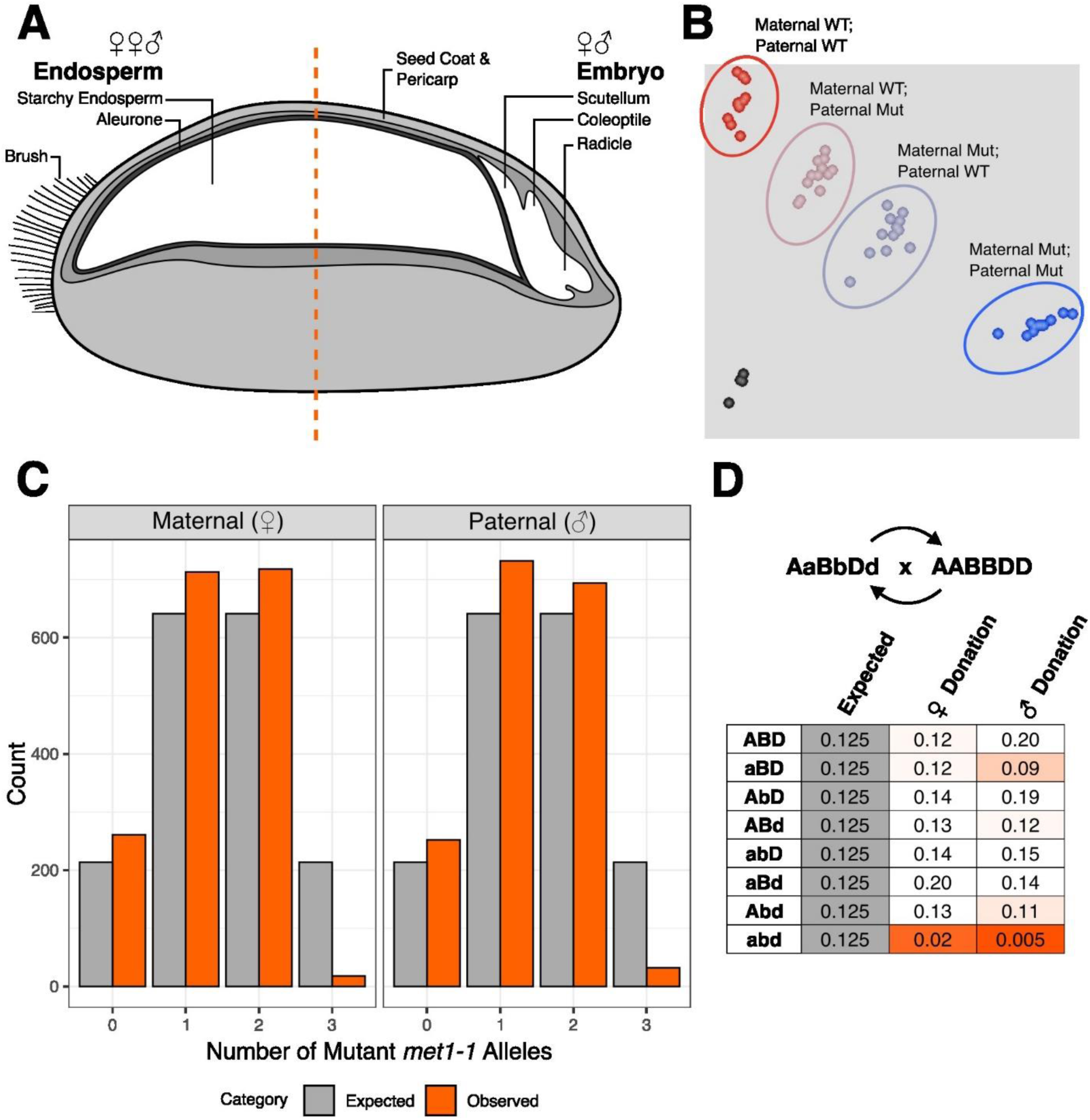
Inheritance of three mutant alleles *met1-1* from the female or male parent (i.e. abd gamete) is rare relative to inheritance of zero, one or two mutant alleles. A) Wheat grains divided transversely to separate two halves of the grain, with one half containing the embryo for future germination and the other half containing largely endosperm for DNA extraction and genotyping. B) KASP genotyping for *Rht13* mutants (*Rht-B13b*) from reciprocal crosses between mutant and wild type parents with known genotypes, showing homozygous wild type (WT) (red cluster), homozygous mutant (blue cluster), heterozygous clusters with a maternally inherited WT allele and paternally inherited mutant allele (brown), a paternally inherited WT allele and maternally inherited mutant allele (grey) and no template control (black). C) Expected (grey) and observed (orange) number of individuals inheriting zero, one, two, or three mutant *met1-1* alleles from the maternal or paternal gamete from a self- pollinated AaBbDd plant. D) Expected and observed proportion of individuals formed from each type of gamete when inherited maternally (♀ donation) or paternally (♂ donation) from an AaBbDd parent when crossed with AABBDD. Orange shades indicate proportions below the expected proportion of 0.125.

We confirmed these observations in an independent segregating F_4_ tetraploid wheat population (332 grains) (Supplementary Figure 3). We observed a significant segregation distortion (χ^2^ p= 3.39x10^-30^) and were unable to identify any homozygous null mutants (aabb). Tetraploid wheat gametes with a complete knock-out of *MET1-1* (i.e. ab gametes) were also strongly underrepresented (generalised linear model with log link, p<2x10^-16^), with reduced maternal and paternal transmission (Supplementary Table 3). In tetraploid wheat we also detected fewer grains which inherited two mutant alleles from the maternal side than from the paternal side (1 vs 18 grains; Supplementary Table 3).

In hexaploid wheat, we found the distribution of genotypes in grains was significantly different from seedling leaf tissue (χ^2^ p=0.00058), with a greater proportion of individuals with five non-functional *MET1-1* alleles observed in the grain population (12 out of 1,710 in the grain population [0.702%] compared to 1 out of 1,352 in the seedling population [0.074%]). This suggests that higher-order *met1-1* mutants may be formed as grains, but not be capable of germination. We hypothesised *met1-1* mutants inheriting three non-functional alleles from the same parent may be associated with decreased grain size. We found that, for grains with a total of three or four mutant *met1-1* alleles, those inheriting three mutant alleles from one parent were smaller by ∼1.5mm than those inheriting the mutant alleles separately from both parents (ordered logistic regression model, p<0.05; Supplementary Figure 4).

To further understand why the recovery of *met1-1* mutants formed from at least one null mutant gamete is reduced, we investigated whether pollen development was affected in the *met1-1* mutants. We measured the number of pollen grains per anther, pollen diameter, pollen viability, and the proportion of pollen grains that were non- or mononucleated for plants from the segregating F_2_ population, including those that should be capable of producing gametes with no functional copies of *MET1-1* (AaBbdd, AabbDd, and aaBbDd). However, no significant differences (at α=0.05) were detected between those with an AaBbdd, AabbDd, or aaBbDd genotype and the WT segregant (AABBDD) for any of these phenotypes (Supplementary Table 4).

To confirm whether inheritance of three mutant *met1-1* alleles from the same parent reduces viability, we performed reciprocal crosses between triple heterozygous (AaBbDd) *met1-1* mutants and Cadenza WT. In total, 454 progeny were genotyped, 235 with AaBbDd as the female parent and 219 with AaBbDd as the male parent. Again, we found that plants inheriting three non-functional *MET1-1* alleles from one parent were significantly underrepresented compared to the expected Mendelian ratio (χ^2^ p<0.001 for both maternal and paternal donation; Figure 2D). The proportion of abd gametes was lower from the paternal side of the reciprocal crosses than the maternal side, opposite to the trend observed in the selfed progeny of a triple heterozygous plant (Figure 2C).

### Loss of four or more functional copies of *MET1-1* results in decreased CG methylation

We carried out whole genome bisulfite sequencing to determine whether a stepwise reduction in the number of functional *MET1-1* alleles was associated with a proportional change in DNA methylation. We observed partial functional redundancy between the *MET1-1* homoeologs; genome-wide CG methylation in the single mutants (aaBBDD, AAbbDD, and AABBdd) was between 87.0% and 87.1% which was very similar to the WT segregant (AABBDD, 87.3% CG methylation). Genome-wide CG methylation was reduced in the double mutants, ranging from 74.0% in the aabbDD mutant, to 79.3% in the aaBBdd mutant, and 81.2% in the AAbbdd mutant. The lowest level of CG methylation was observed in the Aabbdd mutant, with 72.4% (Figure 3A). However, the Aabbdd mutant had the highest CHG (57.4%) and CHH (4.1%) methylation of all the lines tested (Figure 3A).

**Figure 3:**
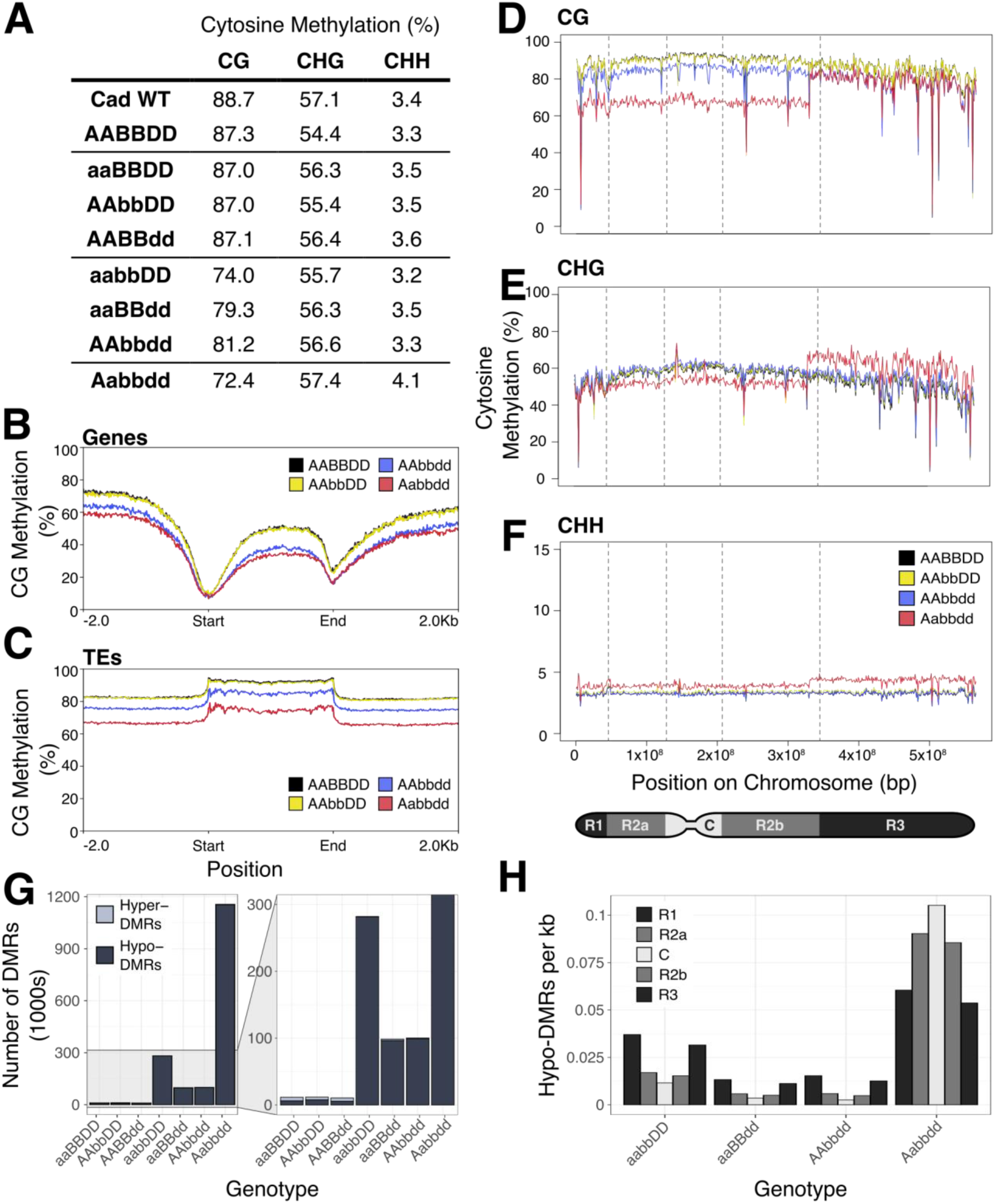
Reduced number of functional copies of *MET1-1* results in loss of CG methylation and the formation of differentially methylated regions (DMRs). A) Genome-wide percentage of cytosines methylated in the CG, CHG, and CHH contexts. B-C) Percentage CG methylation across genes (B) and transposable elements (TEs) (C) on chromosome 5D, including 2 kb up and downstream of the feature. The WT segregant (AABBDD, black), AAbbDD single mutant (yellow), AAbbdd double mutant (blue), and Aabbdd mutant (red) are shown. D-F) Percentage of methylated cytosines in the CG (D), CHG (E), and CHH (F) contexts across chromosome 5D, calculated as an average in 1 Mb bins across the chromosome. The WT segregant (AABBDD, black), AAbbDD single mutant (yellow), AAbbdd double mutant (blue), and Aabbdd mutant (red) are shown. Partitions between the chromosome regions are indicated by the dashed lines. G) Number of differentially methylated regions (DMRs) in 1000s relative to AABBDD in each genotype. Both hypo- methylated (dark grey) and hyper-methylated (light grey) regions are shown. H) Number of hypo-methylated DMRs per kb in each of the chromosome regions – the distal regions R1 and R3 are shown in dark grey, the proximal regions R2a and R2b in medium grey, and the centromeric region C in light grey.

To explore the impact of these global changes in methylation, we examined methylation across genes and transposable elements (TEs), using chromosome 5D as an example. Average CG methylation was decreased across genes and TEs relative to the WT segregant (AABBDD) in both the double mutant (AAbbdd) and the Aabbdd mutant (Fig 3B and C). For both sequence features, the greatest decrease was observed for the Aabbdd mutant, in line with the global changes in methylation. No obvious difference was observed in CHG methylation across genes or TEs, while CHH methylation was highest in the Aabbdd mutant across both genes and TEs (Supplementary Figure 5).

Spatial correlations between different methylation contexts were observed when looking at the distribution of cytosine methylation across each chromosome. Along chromosome 5D, CG methylation is reduced in the Aabbdd mutant compared to the WT segregant, single and double mutants in the R1, R2a, C, and R2b regions. In the R3 region, however, CG methylation in the Aabbdd mutant is comparable to the AAbbdd double mutant (Figure 3D). In this region, there is also an increase in CHG methylation in the Aabbdd mutant, such that it is higher than any of the other lines (Figure 3E), and an increase in CHH methylation (Figure 3F). There is a clear transition point between states in the Aabbdd mutant across all three methylation contexts (Supplementary Figure 6). The simultaneous increase in CG and CHG/CHH methylation is also observed in specific regions of another ten chromosomes, affecting all possible chromosome regions not only R3, and across the whole of chromosomes 3A, 4D and 6B (Supplementary Figures 7-13). These region-specific increases in CG, CHG and CHH methylation were only observed in the Aabbdd mutant, and not in any of the single or double mutants, as shown for chromosome 5D (Supplementary Figure 14).

To further understand the spatial nature of methylation changes in the *met1-1* mutants, we identified 100 bp differentially methylated regions (DMRs) in the CG context in each of the mutants relative to the WT segregant. Substantially more DMRs were detected in the Aabbdd mutant (1,156,551 DMRs), compared to between 98,626 and 282,113 DMRs in the double mutants, and between 10,616 and 11,803 DMRs in the single mutants (Figure 3G). The vast majority of DMRs were hypo-methylated in the *met1-1* mutants – only 0.27% of the DMRs in the Aabbdd mutant were hyper-methylated (Figure 3G). We observed that, in the double mutants, the distal regions R1 and R3 have the greatest density of DMRs, while the centromeric region C has the lowest density (Figure 3H). However, in the Aabbdd mutant the centromeric region has the highest density of DMRs (Figure 3H). In all *met1-1* mutants, the density of hyper-methylated DMRs is lowest in the proximal and centromeric regions, and highest in the distal regions (Supplementary Figure 15).

### Reduced function of *MET1-1* results in stepwise changes in gene expression

To determine the effect of these changes in methylation, we conducted RNA-sequencing and differential expression analysis. The number of differentially expressed genes (DEGs) compared to the WT segregant (FDR adjusted p<0.01, fold change ≥ 2) was calculated for each mutant class (Figure 4A). The Aabbdd mutant had the largest number of DEGs (5,904), while double mutants had on average 79 DEGs, and single mutants had on average 9 DEGs. Generally, there were more up-regulated DEGs than down-regulated DEGs, except for the Aabbdd mutant, with 3,900 genes down-regulated and 2,004 genes up-regulated (Figure 4A). There was also a stronger change in transposable element (TE) expression in the double and Aabbdd mutants compared to single mutants, in which no differentially expressed TEs (DETs) were observed (on average 36 DETs in double mutants and 143 DETs in Aabbdd mutant; FDR adjusted p<0.01, fold change ≥ 2, Figure 4B). These changes in gene and TE expression are in line with the global changes in CG methylation, with mutants maintaining the highest level of CG methylation relative to the WT segregant having the fewest DEGs, and vice versa (Figures 3A, 4A and 4B).

**Figure 4.**
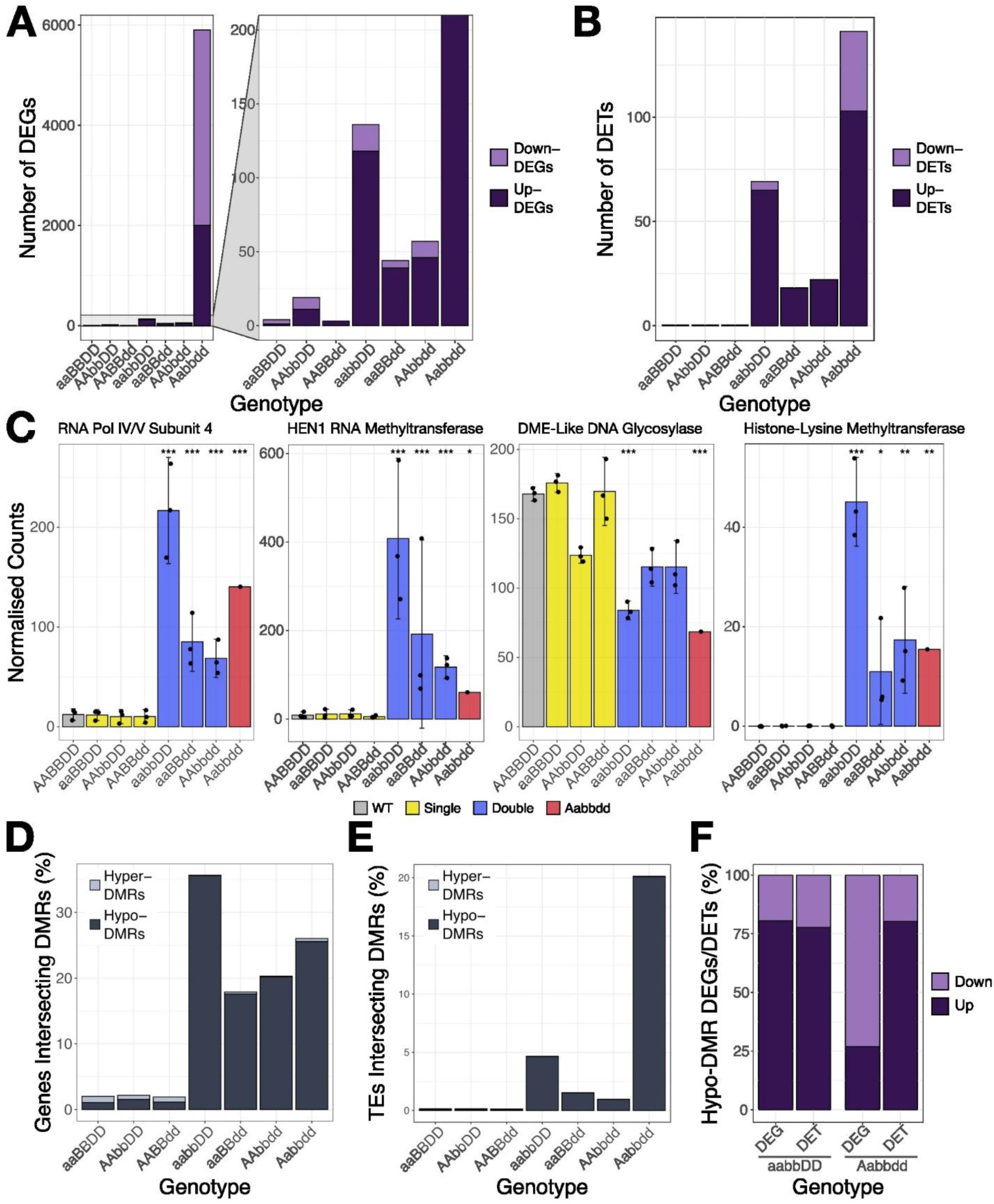
Reduced number of functional copies of *MET1-1* results in transcriptional reprogramming. A) Number of differentially expressed genes (DEGs) relative to AABBDD in each genotype. Both down-regulated (light purple) and up-regulated (dark purple) genes are shown. B) Number of differentially expressed transposable elements (DETs) relative to AABBDD in each genotype (uniquely mapped reads only). Both down-regulated (light purple) and up-regulated (dark purple) transposable elements are shown. C) Normalised transcript counts in each genotype for genes putatively encoding an RNA Polymerase IV/V subunit (*TraesCS6B02G315200*), a HEN1 small RNA 2’-O-methyltransferase (*TraesCS2D02G114600*), a DEMETER (DME)-like DNA glycosylase (*TraesCS3B02G023200*), and a histone-lysine methyltransferase (*TraesCS3B02G503200*). Error bars represent the 95% confidence interval, estimated as 1.96 x standard error. Asterisks show adjusted p-values for differential expression relative to AABBDD; * p<0.05, ** p<0.01, *** p<0.001. D-E) The percentage of genes (D) or transposable elements (E) which intersect at least one DMR in each of the *met1-1* mutants. Hyper-DMRs are shown in light grey and hypo-DMRs are shown in dark grey. F) The percentage of DEGs or DETs associated with a hypo-DMR which are down-regulated (light purple) or up-regulated (dark purple) in the aabbDD and Aabbdd *met1-1* mutants.

To further examine the transcriptional response, we identified genes that were differentially expressed in both the Aabbdd mutant and at least one of the double mutants. A total of 86 DEGs were identified in this way, 76 of which were up-regulated, and 10 of which were down-regulated. Several of these DEGs have putative functions relating to DNA methylation and other epigenetic marks (Figure 4C), suggesting that reduced *MET1-1* function initiates active changes in transcriptional regulation. The gene *TraesCS6B02G315200* is up-regulated in all double mutants and the Aabbdd mutant (adjusted p<0.01) (Figure 4C), and is annotated in the Panther Classification System as RNA Polymerase IV and V Subunit 4 (PTHR21297:SF2). The homoeologous gene *TraesCS6D02G266900* is also up-regulated in all double mutants and the Aabbdd mutant, while its other homoeolog *TraesCS6A02G286200* and tandem duplicated gene copy *TraesCS6B02G15300*, are up-regulated only in the aabbDD and Aabbdd mutants. A gene with the same Panther classification in Arabidopsis, *NRPD4* (*AT4G15950*), is required for RNA-directed DNA methylation (RdDM) (He *et al*., 2009). The gene *TraesCS2D02G114600* was also up-regulated in all double *met1-1* mutants (Figure 4C). This gene encodes a small RNA 2’-O-methyltransferase annotated as HEN1 in the Panther Classification System (PTHR21404). In Arabidopsis, *HEN1* is also implicated in RdDM (Erdmann & Picard, 2020). Furthermore, in both the aabbDD and Aabbdd mutants, *TraesCS3B02G023200*, a DEMETER (DME)-like DNA glycosylase, is significantly down- regulated (Figure 4C). In Arabidopsis, loss-of-function mutations in DME-like DNA glycosylases *DML2* and *DML3* result in increased cytosine methylation in regions with low to intermediate methylation levels in wild type (Ortega-Galisteo *et al*., 2008). Overall, this suggests that reduced *MET1-1* function is buffered by up-regulation of genes that facilitate DNA methylation and down-regulation of genes that remove DNA methylation.

We also observed up-regulation of *TraesCS2B02G503200* in the Aabbdd, aabbDD and AAbbdd mutants (adjusted p<0.01) and the aaBBdd mutants (adjusted p<0.05) (Figure 4C). This gene is predicted to contain a histone-lysine N-methyltransferase SUVR4/SUVR1/SUVR2 domain (IPR025776). Reduced function of *MET1-1* may therefore promote changes in histone methylation that could further alter transcription.

Next, we investigated whether the CG-DMRs identified in the *met1-1* mutants (Figure 3) overlapped genes and TEs. Up to 35.7% of high-confidence genes (IWGSC *et al*., 2018) overlapped with CG-DMRs in the *met1-1* mutants (either within the gene body, or within 1 kb up- or downstream) (Figure 4D). Although the Aabbdd mutant had 4.1-11.7x as many CG-DMRs as the double mutants (Figure 3G), the percentage of genes intersected by at least one CG-DMR (26.0%) was within the range observed in the double mutants (between 17.9% and 35.7%). In line with the overall distribution of DMRs, the majority of DMRs intersecting genes were hypo-methylated in the *met1-1* mutants (Figure 4D).

In contrast to the DMRs intersecting genes, we observed a stark difference between the percentage of TEs overlapped by at least one CG-DMR in the Aabbdd mutant and the double mutants (Figure 4E). In the Aabbdd mutant, 20.1% of non-fragmented TEs (Wicker *et al*., 2018) were intersected by at least one DMR, compared to between 0.95% and 4.7% in the double mutants. This is in line with the greater density of hypo-DMRs in the transposon-dense centromeric region in the Aabbdd mutant (Figure 3H). 87.0% of hypomethylated TEs in the Aabbdd mutant were long terminal repeat retrotransposons (LTR-RTs), compared to between 56.5% and 72.5% in the double mutants (Supplementary Table 5).

To examine whether changes in methylation status may be directly related to changes in gene expression, we quantified the percentage of DEGs which overlapped at least one CG-DMR. We focused on the double and Aabbdd mutants, as only a small number of DEGs were identified in the single mutants (Figure 4A). Between 32.6% and 51.5% of DEGs were associated with a DMR. For each mutant, the proportion of DEGs associated with a DMR is greater than the proportion of genes associated with a DMR, suggesting that changes in DNA methylation may be directly responsible for the change in expression of a subset of DEGs. In the aabbDD double mutant, 80.6% of hypomethylated DEGs increased in expression relative to the WT segregant (Figure 4F). However, in the Aabbdd mutant, only 26.9% of hypomethylated DEGs were up-regulated (Figure 4F). This could reflect the overall greater proportion of down-regulated genes in the Aabbdd mutant compared to the double mutants (Figure 4A). The majority of DETs associated with a hypo-DMR were up-regulated in both the aabbDD and Aabbdd mutants (77.8% in aabbDD and 80.4% in Aabbdd) (Figure 4F). The same trend was observed for the other double mutants (Supplementary Figure 16).

### Heritable phenotypic changes were identified in the *MET1-1* segregating F2 population

When growing the C0451xC1292xC2028 F_2_ population we identified one plant (AaBbdd genotype) that flowered 23 days later than any other plant and had lax ears (thin ears composed of many spikelets). Offspring from this late-flowering individual segregated into normal flowering types with normal ears (<10 days relative to earliest flowering plant), and late-flowering types with lax ears (≥10 days relative to earliest flowering plant) (Figure 5A- C). The numbers of segregant plants in the late and normal flowering categories did not significantly differ from a 3:1 ratio (χ^2^ = 0.214, p-value = 0.644), suggesting that a single dominant gene controls the phenotype. The late flowering phenotype was independent of *MET1-1* genotype (Figure 5D). The late flowering plants were also significantly taller than normal flowering plants (FDR adjusted p-value = 8.2x10^-15^; Figure 5A, E). There was no significant difference in spike length between the late and normal flowering segregants (FDR adjusted p-value = 0.22, Figure 5F), however late flowering segregants had a higher number of fully formed spikelets per spike than normal flowering segregants (mean 25.3 vs 18.6; FDR adjusted p-value = 5.5x10^-9^, Figure 5B and G). These data show that the tall, late flowering phenotype is heritable and is independent of *MET1-1* genotype, suggesting that it is controlled by a single stable allele generated in the mutant background.

**Figure 5.**
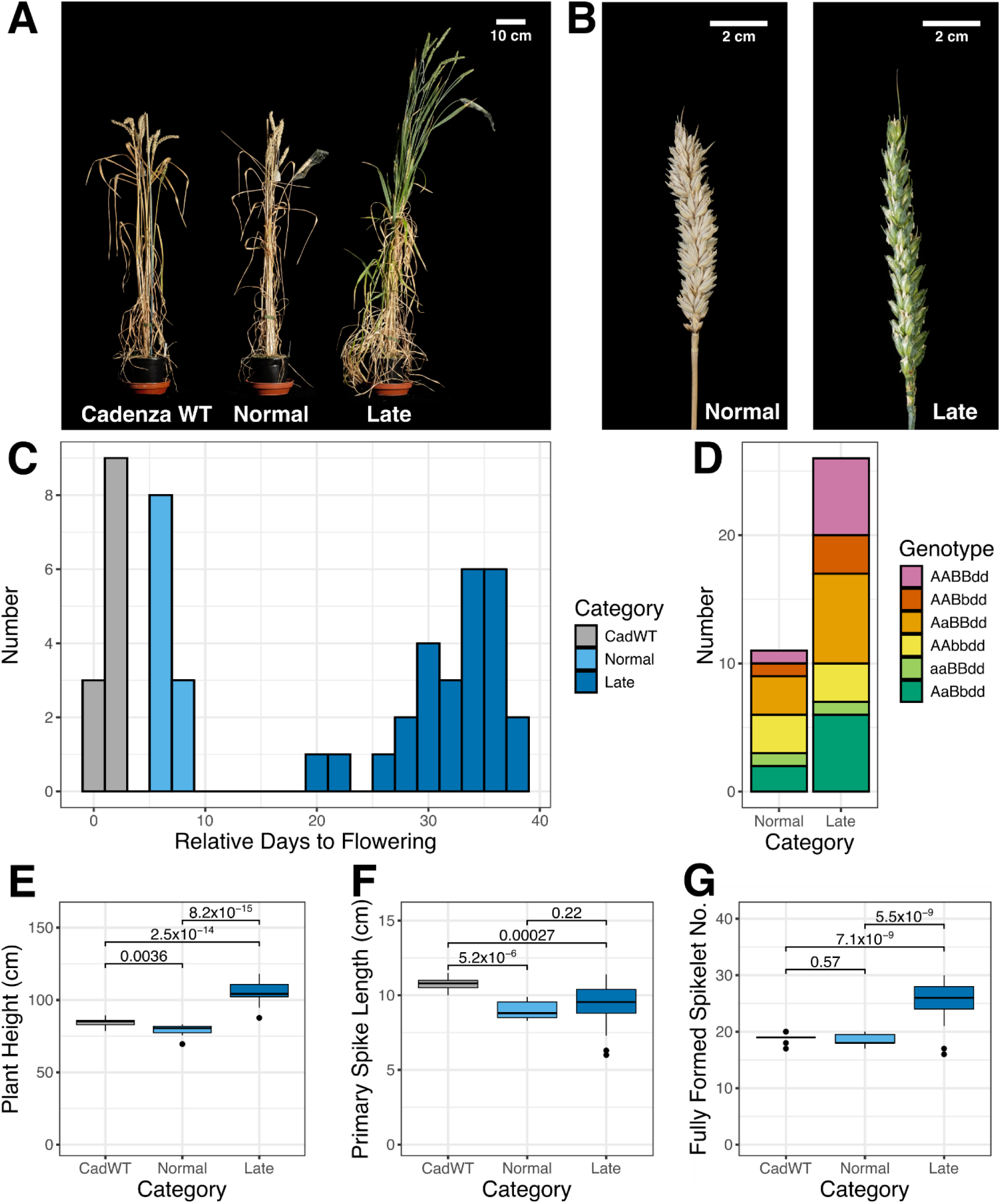
The late flowering phenotype in the C0451xC1292xC2028 *MET1-1* population was independent of *MET1-1* genotype and was accompanied by increased plant height and number of fully formed spikelets. A) Phenotypes of Cadenza WT and normal and late flowering F_3_ segregant plants. B) Normal and late flowering segregant plants showing the increased number of fully formed spikelets in the late flowering plant. C) Histogram showing the days to flowering (full emergence of primary spike) of the different flowering time categories relative to the first plant. D) *MET1-1* genotypes of late and normal flowering plants. Boxplots of plant height (E), primary spike length (F) and number of fully formed spikelets (G) for each of the phenotype categories. For (E) to (G), the means were compared by a t-test and the False Discovery Rate (FDR) adjusted p-value calculated is shown.

A second phenotype was observed in the same segregating F_2_ population (C0451xC1292xC2028): one plant (AAbbdd genotype) exhibited an awned phenotype, despite all three parents being awnless. We generated a segregating population by backcrossing the awned plant with Cadenza WT and selfing the F_1_ generation, which showed an intermediate (awnletted) phenotype. Amongst 215 F_2_ plants the segregation ratio of awnless:awnletted:awned individuals did not significantly deviate from a single co-dominant gene 1:2:1 Mendelian ratio (χ^2^ = 1.13, p-value = 0.568) (Figure 6A and B). The awnletted individual had longer awns at the top of the ear, similar to the effect of the *Tipped1* (*B1*) awn suppressor gene (Huang *et al*., 2020) and we hypothesised that this was the causal gene controlling the phenotype. We used published primer sequences to screen the *B1* region in the F_2_ population (Figure 6C) (Huang *et al*., 2020). PCR products could be obtained for awnless and awnletted plants, but not for awned plants either across the gene body or in the surrounding 28.5 Mbp (Figure 6C and D). Together, these results indicate that the presence of awns in our population is due to a heritable deletion >28.5Mbp which includes *B1*.

**Figure 6.**
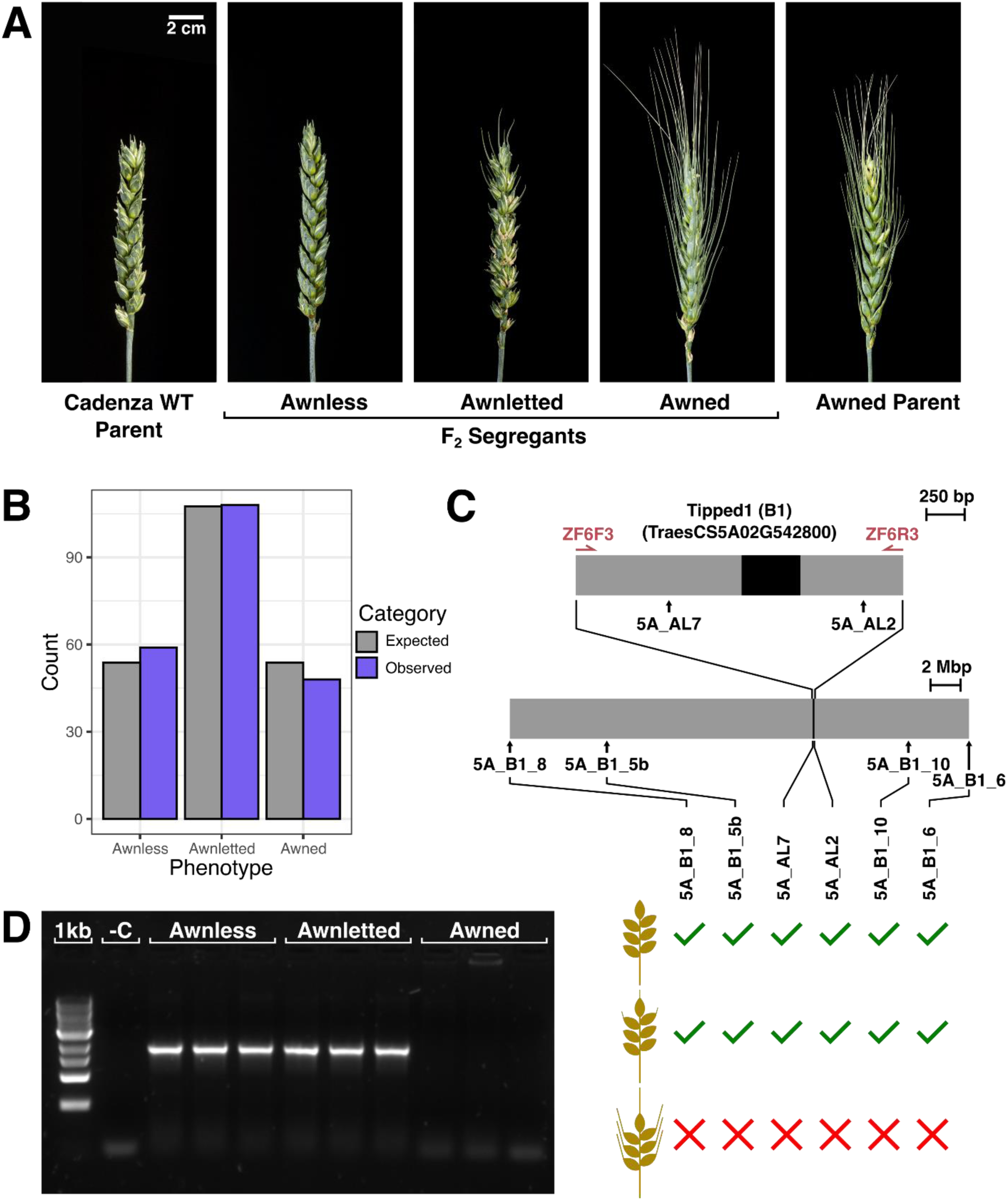
The observed awned phenotype in the C0451xC1292xC2028 *MET1-1* population followed a single gene semi dominant mendelian inheritance pattern, which was caused by a deletion encompassing *Tipped1* (*B1*). A) Phenotypes of the awnless Cadenza WT parent and spontaneous awned individual which were crossed together to generate the F_1_ generation. The F_1_ was self-pollinated to generate the segregating F_2_ population which displayed awnless, awnletted and awned phenotypes. B) Expected (grey) and observed (purple) number of individuals which were awnless, awnletted and awned in the segregating F_2_ population. C) Diagram of the *B1* gene (black) and surrounding region (grey) with KASP marker binding sites (black arrows) and PCR primer binding sites (maroon half arrows). Schematic summarising successful (green tick) and unsuccessful (red cross) amplification of KASP marker sets for each phenotype class of plant (awned, awnletted and awnless). D) Agarose gel image showing PCR product of the ZF6F3 and ZF6R3 primer set (expected product ≈2kb) from 3 individual F_2_ segregant plants of each phenotype class and 1kb DNA ladder (1kb) and no DNA (-C) controls.

## Discussion

### Complete loss of *MET1-1* is lethal in wheat, but single and double mutants can be recovered

Despite genotyping over 3,000 progeny from two segregating *MET1-1* populations, we were unable to recover any null *met1-1* mutants in hexaploid wheat (Figure 1F, Supplementary Table 1). Similarly, we were unable to recover any null *met1-1* mutants in tetraploid wheat (Supplementary Table 3). This indicates that complete loss of *MET1-1* is lethal in wheat, consistent with other monocotyledonous crop species with large genomes such as maize and barley (Li *et al*., 2014; Schreiber *et al*., 2019).

However, unlike maize and barley, wheat is a polyploid, allowing intermediate mutants to be recovered. We recovered all possible homozygous single and double *met1-1* mutants from the segregating *MET1-1* populations suggesting that the function of *MET1-1* homoeologs in wheat is mostly redundant. We also recovered a single Aabbdd mutant, in which five out of six copies of *MET1-1* are non-functional (Figure 1D). We found that *MET1-1* has a threshold for dosage-sensitivity: while the single mutants had no decrease in CG methylation compared to the WT segregant, the double and Aabbdd mutants showed altered patterns of CG methylation and gene expression (Figure 3 and 4). The double mutant classes showed variability in the degree of CG methylation loss and changes to gene expression, with the aabbDD mutants showing the greatest change compared to the AAbbdd and aaBBdd double mutants (Figure 3 and 4). This suggests that *MET1-D1* is less effective at compensating for the loss of the other two homoeologs than *MET1-A1* or *MET1-B1*. Furthermore, the double mutants did not show any growth defects, despite the aabbDD mutants having a similar degree of demethylation to the Aabbdd mutant (74.0% CG methylation and 72.4% CG methylation respectively). Together these features suggests that double mutants with four mutant alleles may allow the generation of epialleles which could be utilised in epigenetic breeding (Figure 7).

**Figure 7.**
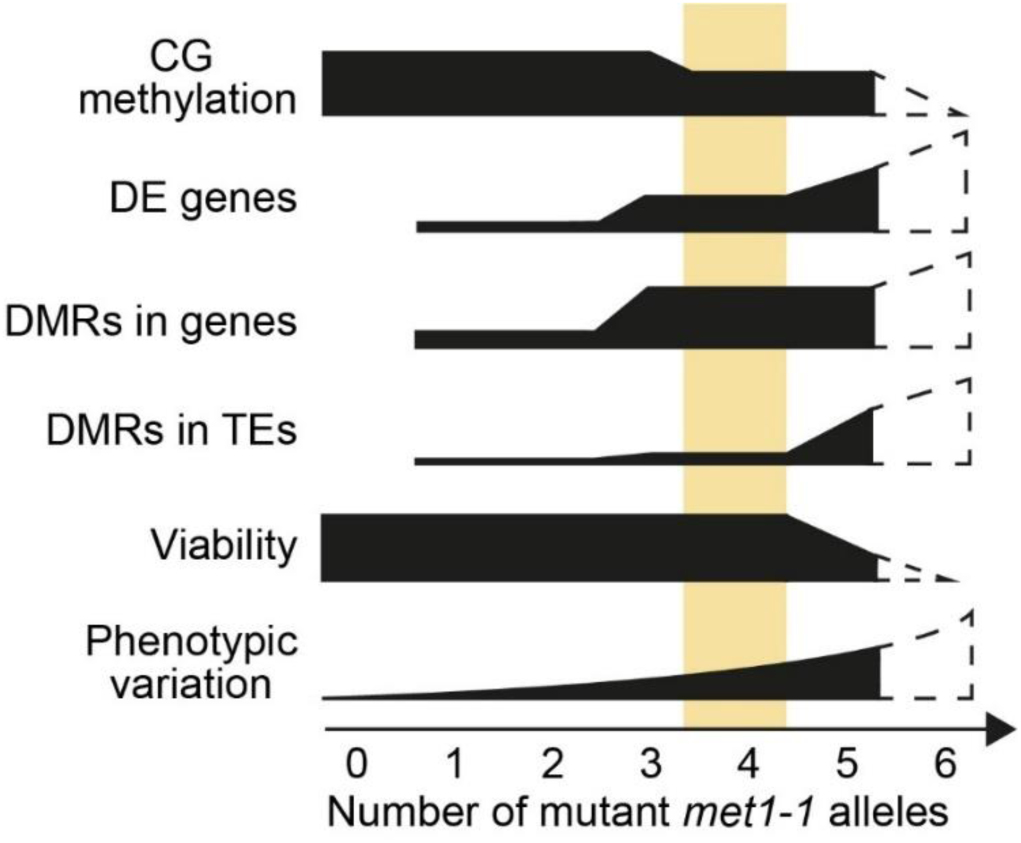
Increasing the number of mutant *met1-1* alleles in hexaploid wheat has quantitative yet variable effects on CG methylation, differentially expressed (DE) genes, differentially methylated regions (DMRs) in genes and in transposable elements (TEs), viability and phenotypic variation. Dashed regions indicate hypothetical data for a complete null mutant. The region highlighted in yellow represents an optimum where CG methylation is substantially reduced and viability is retained, made possible by partial redundancy between homoeologs.

We observed novel, heritable phenotypic variation in one of the segregating *MET1-1* populations (Figure 5 and 6). These phenotypes (delayed flowering and the development of awns) were both identified in plants that had four out of six non-functional copies of *MET1-1*, indicating that a null *met1-1* mutant is not necessary to generate phenotypic variation. Loss of DNA methylation can generate novel variation in multiple ways. Firstly, epialleles may be generated, resulting in altered patterns of gene expression. Secondly, hypomethylation of transposable elements (TEs) is associated with their transcription and remobilisation, leading to abnormal developmental phenotypes (Miura *et al*., 2001; Mirouze *et al*., 2009; Reinders *et al*., 2009; Tsukahara *et al*., 2009). We found that one fifth of TEs were associated with at least one DMR in the Aabbdd mutant, and 45 of these TEs had up-regulated expression, potentially resulting in their activation. This may explain the stunted growth of the Aabbdd mutant, and the severely reduced viability of higher-order *met1-1* mutants. Genome instability caused by loss of DNA methylation can also cause structural variation (Zhang *et al*., 2023), which we observed in the spontaneous awned *met1-1* mutant that had a substantial deletion (Figure 6).

A similar large deletion was found in wheat treated with the demethylation agent zebularine; the deletion encompassed *FT-B1* and was associated with an increase in spikelet number and a delay in flowering (Finnegan *et al*., 2019).

### Lethal effect of null *met1-1* mutants associated with mutant (abd) gametes

We observed a clear segregation distortion when genotyping selfed seed from a triple heterozygous (AaBbDd) *MET1-1* parent, with underrepresentation of higher-level mutants (5/6 and 6/6 non-functional copies). We attributed this distortion to a reduced transmission of gametes with no functional copy of *MET1-1* (abd gametes) when donated maternally or paternally. In reciprocal crosses, abd gametes transmitted paternally were rarer than abd gametes transmitted maternally, whereas the opposite trend was observed using selfed heterozygous (AaBbDd) mutants. This discrepancy may be explained by post-fertilisation effects of *met1-1* being made more severe in a scenario where mutant alleles may be present in both gametes, as for the selfed heterozygous plants. This is consistent with work in Arabidopsis that found that selfed heterozygous *met1* mutants had higher rates of embryo abortion than wild type x homozygous *met1* mutant reciprocal crosses (Xiao *et al*., 2006).

In Arabidopsis, transmission of mutant *met1* alleles has also been shown to be distorted in the gametes. However, in contrast to our results where both maternal and paternal transmission was affected, only the paternal transmission of *met1* was reduced to 36.8% of the expected rate in Arabidopsis while maternal transmission was not reduced (Liang *et al*., 2022). This single parental bias in *met1* transmission was, in part, due to impaired pollen development resulting in abnormal numbers of nuclei in the pollen grains. In our work we found no change to any pollen phenotype measured in the *met1-1* mutant plants compared to WT segregant plants, suggesting that instead the barrier to the production of null mutants may occur during fertilisation and/or embryogenesis. Post-zygotic effects are also likely important since we found over 9x more offspring with five non-functional *MET1* alleles in grains than in leaves of seedlings. Reduced germination may be due to abnormal endosperm development as observed in Arabidopsis *met1* mutants (Jullien *et al*., 2006). This would also be consistent with the smaller seed sizes we observed in wheat mutants inheriting *met1-1* alleles through abd gametes, although the effect was not as extreme as observed in rice (Hu *et al*., 2014; Yamauchi *et al*., 2014), nor was it exclusively paternal as in Arabidopsis (Xiao *et al*., 2006).

### Reduced function of *MET1-1* is buffered by activation of other epigenetic pathways

Loss of five out of six functional copies of *MET1-1* resulted in a significant decrease in CG methylation, but an increase in CHG and CHH methylation (Figure 3), suggesting that other methylation pathways are activated upon reduction of *MET1-1* activity. A similar increase in CHG and CHH methylation was observed in tomato *met1* mutants (Yang *et al*., 2019) and in rice *MET1/met1* heterozygotes, but not the homozygous *met1* mutant (Hu *et al*., 2014). In Arabidopsis, CHG methylation was increased in *met1* mutants, while CHH methylation was decreased (Srikant *et al*., 2022). In wheat, the increases in CHG and CHH methylation are found in extremely well-defined regions where CG methylation is maintained to at least the level of the AAbbdd double mutant (Figure 3, Supplementary Figures 5-12). Future studies are required to determine whether these regions feature only in species with large genomes, making them undetectable until now.

The compensatory increase in CHG and CHH methylation we observed may be promoted by transcriptional up-regulation of RNA-directed DNA methylation (RdDM). We found increased expression of genes involved in generating short interfering RNAs (siRNAs) which direct the RdDM machinery to loci to be methylated (He *et al*., 2009; Erdmann & Picard, 2020); these include RNA Polymerase IV/V Subunit 4 and the HEN1 RNA methyltransferase in the double and Aabbdd *met1-1* mutants (Figure 4C). We also found down-regulation of a DME-like DNA glycosylase in the mutants with the largest reduction in CG methylation (aabbDD and Aabbdd) (Figure 4C). Down-regulation of DNA glycosylases has also been observed in *met1* mutants in Arabidopsis and rice (Mathieu *et al*., 2007; Hu *et al*., 2014).

Transcriptional down-regulation of the Arabidopsis *ROS1* DNA glycosylase is caused by loss of methylation in its promoter at the methylation monitoring sequence (MEMS) (Lei *et al*., 2015). We found that the DME-like DNA glycosylase down-regulated in the aabbDD and Aabbdd *met1-1* mutants overlapped a hypo-DMR, suggesting that a similar MEMS mechanism may regulate the expression of this DNA glycosylase, allowing the plant to sense changes in the DNA methylation status and compensate for them.

Compensation for reduced CG methylation can also occur through changes in histone modifications. We observed transcriptional up-regulation of a SUVR4/SUVR1/SUVR2 histone lysine methyltransferase in the double and Aabbdd *met1-1* mutants (Figure 4C). SUVR4/SUVR1/SUVR2 histone lysine methyltransferases have a preference for H3K9 methylation (Thorstensen *et al*., 2006), suggesting that this epigenetic mark may also be altered upon reduction of CG methylation in wheat, as has been reported in Arabidopsis *met1* mutants (Tariq *et al*., 2003; Mathieu *et al*., 2007).

In summary, we leveraged partial redundancy between homoeologs in polyploid wheat to recover *met1-1* mutants. Similar to other plants with large genomes, complete null mutants were lethal. However, wheat plants with four null alleles (i.e. retaining two functional alleles) develop normally, have substantially reduced CG methylation and occasionally exhibit novel heritable phenotypes. Generating partial mutants may also be effective in other polyploid species to manipulate CG methylation without lethal consequences, opening new avenues to understand the role of CG methylation in polyploid species and generate epialleles for crop breeding.

## Methods

### Plant materials and growth

Exome capture sequenced EMS mutagenized populations of hexaploid (*Triticum aestivum* cv. Cadenza) and tetraploid wheat (*Triticum turgidum* cv. Kronos) were screened for premature termination codon mutations in *MET1-1* (*TraesCS2A02G235900*, *TraesCS2B02G260800* and *TraesCS2D02G241800*) predicted to eliminate C5 Cytosine methyltransferase domain activity (Figure 1C) (Krasileva *et al*., 2017).

Two independent Cadenza *MET1-1* populations were developed. The C0465xC0884 population was developed by crossing the double *met1-1* mutant Cadenza0465 with Cadenza0884 to generate a triple heterozygous F_1_, from which a segregating F_2_ population was produced (Supplementary Figure 2). The C0451xC2028xC1292 population was developed by crossing Cadenza0451 with Cadenza2028 and Cadenza2028 with Cadenza1292. The F_1_ progeny were crossed to produce a triple heterozygous F_1_ which was self-pollinated to generate a segregating F_2_ population (Supplementary Figure 2).

The Kronos *MET1-1* population was produced by crossing Kronos3085 with Kronos0809, giving double heterozygous F_1_ progeny, which were selfed to generate a segregating F_2_ population. From the F_2_ population, seeds from double heterozygous plants were sown to give a segregating F_3_ generation. F_4_ seeds from double heterozygous F_3_ plants were used for seed genotyping (Supplementary Figure 3).

Seeds were germinated in petri dishes at 4°C for 48 hours, followed by room temperature for 48 hours before sowing into 96-cell trays containing John Innes F_2_ starter + Grit (90% peat, 10% grit, 4kg m^-3^ dolomitic limestone, 1.2kg m^-3^ osmocote start). Selected plants were potted on into 1L pots containing John Innes Cereal Mix (65% peat, 25% loam, 10% grit, 3kg m^-3^ dolomitic limestone, 1.3kg m^-3^ pg mix, 3kg m^-3^ osmocote exact). Plants were grown in the John Innes Centre glasshouses (Norwich, UK) with supplementary lighting and heating as required for minimum 16h light with 16°C day and 14°C night.

### Genotyping

#### Leaf

At the second leaf stage (Zadoks growth stage 12), DNA was extracted from 2cm leaf samples following the protocol from www.wheat-training.com adapted from Pallotta *et al*. (2003). Genotyping was carried out using KASP markers (Supplementary Table 6) and PACE mix, according to the manufacturer’s instructions. Kluster-Caller (v3.4.1.36) software was used for data analysis. We developed a CAPS marker for C1292 (Supplementary Table 6), which was amplified using Taq DNA Polymerase (New England Biolabs, 94°C for 2:00, 40 cycles of 94°C for 0:15, 50°C for 0:25, 72°C for 1:00 ending with 2:30 final extension at 72°C). The product was digested by restriction enzyme PstI-HF (New England Biolabs, 37°C, 1 hour). Homozygous *MET1-1* plants produced 3 digestion products (94bp, 157bp, 249bp) and homozygous *met1-1* mutants produced 2 digestion products (94bp and 406bp).

#### Grain

Grains were surface sterilised and cut transversely. The half of the grain containing the embryo was germinated and the other half (containing endosperm) was used for genotyping. Full details are available at dx.doi.org/10.17504/protocols.io.6qpvr84xplmk/v1. Parental donation of the *met1-1* allele was determined using KASP genotyping, where the usual single heterozygous cluster was split into two, corresponding to maternal or paternal donation of the mutant *met1-1* allele. This method was validated using reciprocal crosses between *Rht-B13b* (semi-dwarf) and wild type plants using KASP markers from (Borrill *et al*., 2022).

### Phenotyping

#### Grain size measurements

Grains were graded into size categories using 3D printed sieves with apertures from 1.0mm – 3.0mm increasing by 0.5mm increments The sieve design is available at https://www.printables.com/model/890045-grain-sieve. Following grain size grading, grain genotyping was carried out. The effect of total number of mutant *met1-1* alleles and inheritance of an abd gamete on grain size was tested using an ordered logistic regression model with the MASS package in R (v4.2.2).

#### Pollen Assays

Pollen grains per anther and pollen grain diameter were analysed using the Multisizer 4e Coulter Counter (Beckman Coulter, California, USA), based on the method described by (Alabdullah *et al*., 2021). Pollen viability was analysed using the Ampha Z32 Pollen Analyzer (Amphasys, Lucerne, Switzerland). To determine the number of nuclei per pollen grain, pollen grains were stained with DAPI (4’,6-diamidine-2-phenylindole) and imaged. Full method details can be found in the Supplementary Information.

#### Flowering Time

A late-flowering plant from the C0451xC2028xC1292 F_2_ population was selfed. The progeny were genotyped and the date of complete emergence of the primary spike (Zadok’s growth stage Z59) recorded. Plant height and the length of the primary spike were measured. The number of fully formed spikelets on the primary spike were counted.

#### Awn Development

An awned plant from the C0451xC2028xC1292 F_2_ population was crossed to Cadenza WT. BC_1_F_1_ plants were grown and selfed.

215 segregating BC_1_F_2_ plants were grown and scored for the presence of awns as previously described (Huang *et al*., 2020). Fully awned individuals had awns >1cm throughout the spike whereas awnletted plants had awns <1cm in the mid and basal portions of the spike while the length of awns at the apex reached up to 3cm. We scored any plants that had awns <1cm across the entire spike as awnless. We investigated the *Tipped1* (*B1*) gene (*TraesCS5A02G542800*), a characterised awn suppression gene, using published PCR and KASP markers (Huang *et al*., 2020).

### Reciprocal crossing

To test the transmissibility of *met1-1* mutant alleles in each of the parental gametes, we made reciprocal crosses between Cadenza WT and triple heterozygous plants (AaBbDd) from the C0465xC0884 population. The seeds of these crosses were harvested and grown. Once the seedlings reached the two-leaf stage (Zadoks growth stage Z12) leaf genotyping was carried out.

### Whole Genome Bisulfite Sequencing

We collected leaf samples from the fourth leaf at Zadok’s growth stage Z15.6/23, or from the sixth leaf at Z16 for the Aabbdd mutant. Samples were frozen in liquid nitrogen and stored at -70°C. DNA was extracted using the DNeasy Plant Mini Kit (Qiagen, Hilden, Germany) from three biological replicates per genotype, excluding the Aabbdd genotype, of which only one plant was recovered. DNA concentration was measured using an Invitrogen Qubit 4 fluorometer (Thermo Fisher Scientific, Massachusetts, USA) and the three replicates per genotype were pooled in equimolar ratios. Library preparation, bisulfite conversion, and sequencing by Illumina NovaSeq 6000 were performed by BMKgene (Münster, Germany), to obtain an average of 2.62 billion 150 bp paired-end reads per sample. Bisulfite conversion rates were calculated by BMKgene using the Bismark software (Krueger & Andrews, 2011), and rates between 99.52% and 99.70% were obtained. Fastp (Chen *et al*., 2018) was used to remove adapter sequences, low quality reads, and polyG sequences. The Bismark Bisulfite Mapper (Krueger & Andrews, 2011) was used to process the sequencing data. Reads were aligned to a bisulfite-converted version of the IWGSC RefSeq v1.0 reference genome (IWGSC *et al*., 2018). The average mapping efficiency was 40.49%, resulting in average coverage of 11X. Methylation calls in all sequence contexts (CpG, CHG, CHH) were extracted. Average cytosine methylation across genes and transposable elements was plotted using deepTools (Ramírez *et al*., 2016) using GTF files generated using gffread (Pertea & Pertea, 2020) from previously published gff3 files (IWGSC *et al*., 2018; Wicker *et al*., 2018). Only cytosines with at least 4 reads were included in the analysis. Methylation profiles across chromosomes were calculated using computeMethylationProfile from DMRCaller (Catoni *et al*., 2018), with bins of 1 Mb. Partitions between chromosome segments (R1, R2a, C, R2b, and R3) were assigned based on the segments defined in (IWGSC *et al*., 2018), available at https://opendata.earlham.ac.uk/wheat/under_license/toronto/Ramirez-Gonzalez_etal_2018-06025-Transcriptome-Landscape/data/TablesForExploration/. Differentially methylated regions (DMRs) in the CG context were called for each chromosome separately using computeDMRs from DMRCaller (Catoni *et al*., 2018), specifying the method as ‘bins’, binSize as 100 bp, pValueThreshold as 0.01, minCytosinesCount as 10, minProportionDifference as 0.4, minGap as 0, and minReadsPerCytosine as 4. DMRs associated with genomic features were identified using bedtools intersect (Quinlan & Hall, 2010) to find overlapping genomic coordinates between previously annotated high-confidence genes (including 1 kb up- and downstream of the coding sequence) (IWGSC *et al*., 2018) and transposable elements (Wicker *et al*., 2018).

### RNA-Sequencing

We collected leaf samples from the fourth leaf at Zadok’s growth stage Z15.6/23, or from the sixth leaf at Z16 for the Aabbdd mutant. The samples were frozen in liquid nitrogen and stored at -70°C. RNA was extracted using trizol-chloroform and the RNA Clean & Concentrator Kit (Zymo Research). RNA was extracted from three biological replicates per genotype, excluding the Aabbdd genotype. Library preparation and sequencing by Illumina NovaSeq 6000 was performed by BMKgene (Münster, Germany), to obtain 150 bp paired-end reads. Fastp (Chen *et al*., 2018) was used to remove adapter sequences, low quality reads, and polyG sequences. Reads were pseudoaligned to the IWGSC RefSeq v1.1 reference transcriptome (IWGSC *et al*., 2018) and quantified using kallisto (Bray *et al*., 2016). The output from kallisto was imported into R (v4.3.2) using tximport (Soneson *et al*., 2015).

Differentially expressed genes between genotypes were identified using DESeq2, using an adjusted p-value threshold of <0.01 and ≥ 2 fold change (Love *et al*., 2014). For TE expression, reads were mapped to the IWGSC reference sequence using HISAT2 (Pertea *et al*., 2016) and reads uniquely mapping to complete TE positions (Wicker *et al*., 2018) were counted using HTSeq (Anders *et al*., 2015), with default parameters. Differential expression analysis was carried out as for genes.

Gene expression data across tissues was downloaded as tpm values from https://bar.utoronto.ca/efp_wheat/cgi-bin/efpWeb.cgi (Choulet *et al*., 2014; Borrill *et al*., 2016; Ramírez-González *et al*., 2018).

### Data availability

Raw reads from whole genome bisulfite sequencing have been deposited in the European Nucletoide Archive under project PRJEB77426 and reads from RNA-sequencing have been deposited in project PRJEB77425.

All scripts used for whole genome bisulfite sequencing and RNA-sequencing analyses can be found on GitHub at https://github.com/Borrill-Lab/Wheat_MET1/.

## Author contributions

MC and PB conceived the study. SB, DD and PB designed the research. JS and SB generated the mutant populations. SB and DD performed most experiments and statistical analysis. DD carried out the data analysis for the RNA-seq and whole genome bisulfite sequencing. JC assisted with genotyping and phenotyping of awn development. GDS helped with genotyping and carried out pollen phenotyping experiments. DD analysed the pollen phenotyping data. SB, DD and PB created the figures. SB, DD and PB wrote the manuscript with input from all authors. All authors have read and approved the manuscript.

## Acknowledgements

We thank Abdulkader Alabdullah for advice on pollen phenotyping, Marek Glombik for advice on genomic analysis and James Brown for advice on statistical analysis of grain genotyping data. We thank Phil Robinson from Scientific Photography at JIC for taking high-quality images of plants and spikes. This research was supported by the NBI Research Computing group through HPC resources.

