## Supplementary Figures, Tables and Methods for "Partial redundancy buffers deleterious effects of mutating *DNA methyltransferase 1-1* (*MET1-1*) in polyploid wheat"

### Supplementary Figures and Tables for "Partial redundancy buffers deleterious effects of mutating *DNA methyltransferase 1-1* (*MET1-1*) in polyploid wheat"

Samuel Burrows[1]*, Delfi Dorussen[1]*, Joseph Crudgington[1], Giorgia Di Santolo[1], James Simmonds[1], Marco Catoni[2], Philippa Borrill[1†]

[1] Department of Crop Genetics, John Innes Centre, Norwich Research Park, Norwich, NR4 7UH, United Kingdom.

[2] School of Biosciences, University of Birmingham, Birmingham, B15 2TT, United Kingdom.

* These authors contributed equally

#### Supplementary Figures

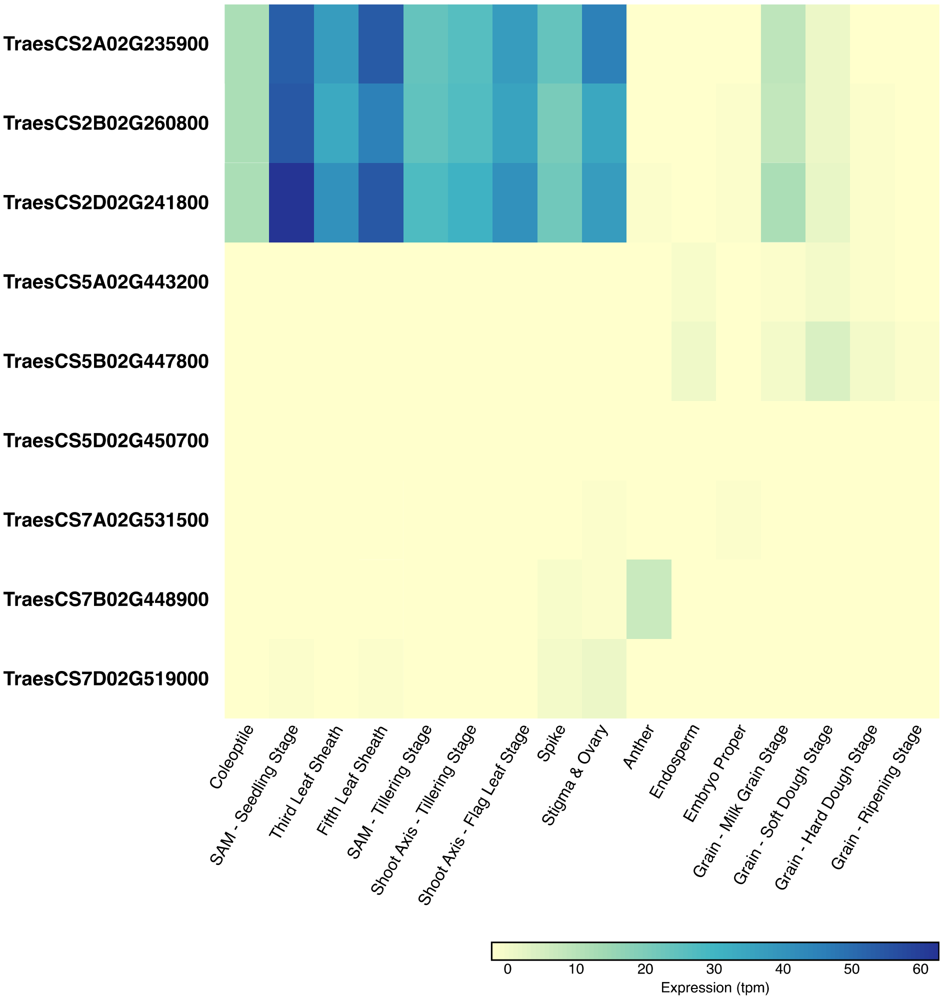


**Supplementary Figure 1.** Expression profiles of Ta*MET1* genes in the cultivar Chinese Spring showing that the chromosome 2 homoeologue group formed of *TraesCS2A02G235900*, *TraesCS2B02G260800* and *TraesCSD02G241800* is the most highly expressed compared to homoeologue groups on chromosome 5 (*TraesCS5A02G443200*, *TraesCS5B02G447800*, *TraesCS5D02G450700*) and chromosome 7 (*TraesCS7A02G531500*, *TraesCS7B02G448900*, *TraesCS7D02G519000*)

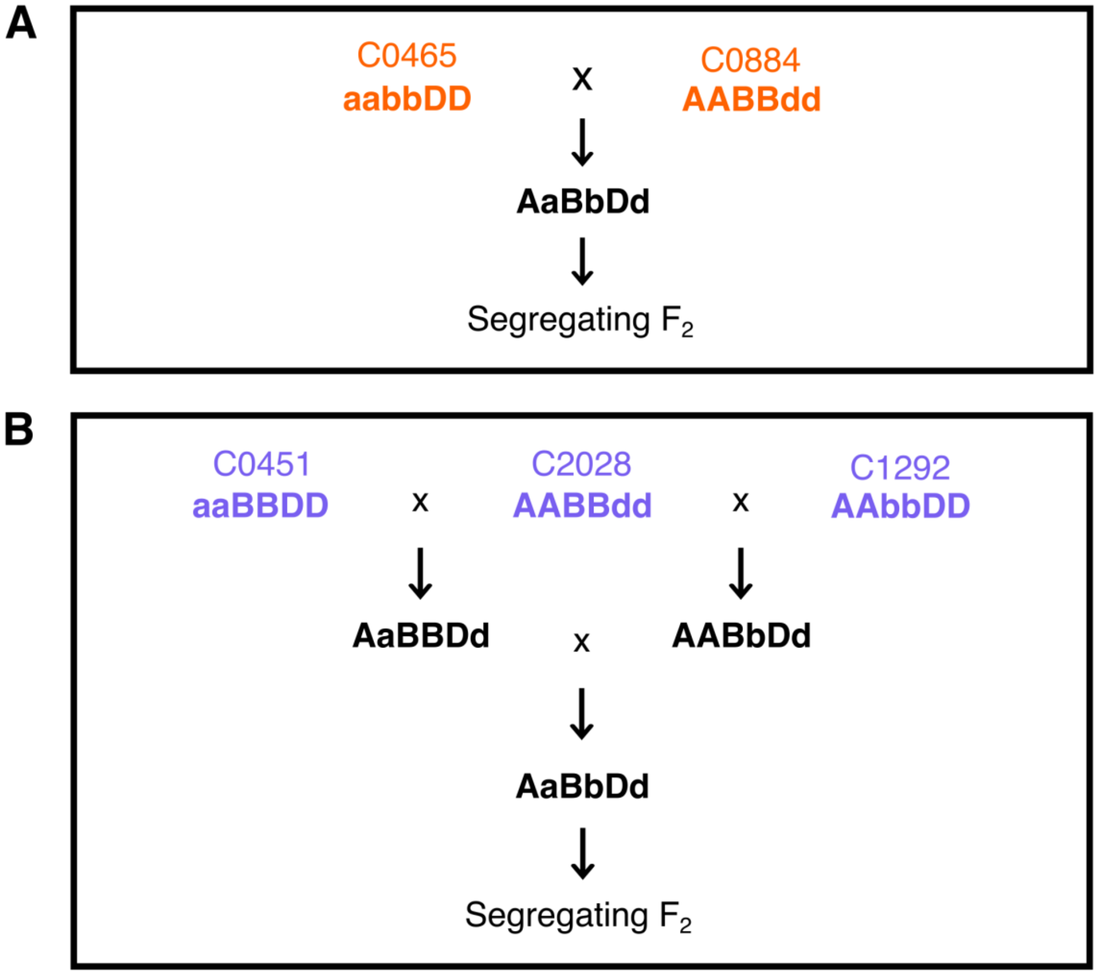


**Supplementary Figure 2.** Crossing schematic showing the generation of the hexaploid *MET1-1* populations. A) Generation of the C0465xC0884 population, formed by crossing 2 TILLING mutant lines (C0465 and C0884) to generate a segregating $F_2$ population. B) Generation of the C0451xC2028xC1292 population, formed by crossing 3 TILLING mutant lines (C0451, C2028, C1292) to generate a segregating $F_2$ population.

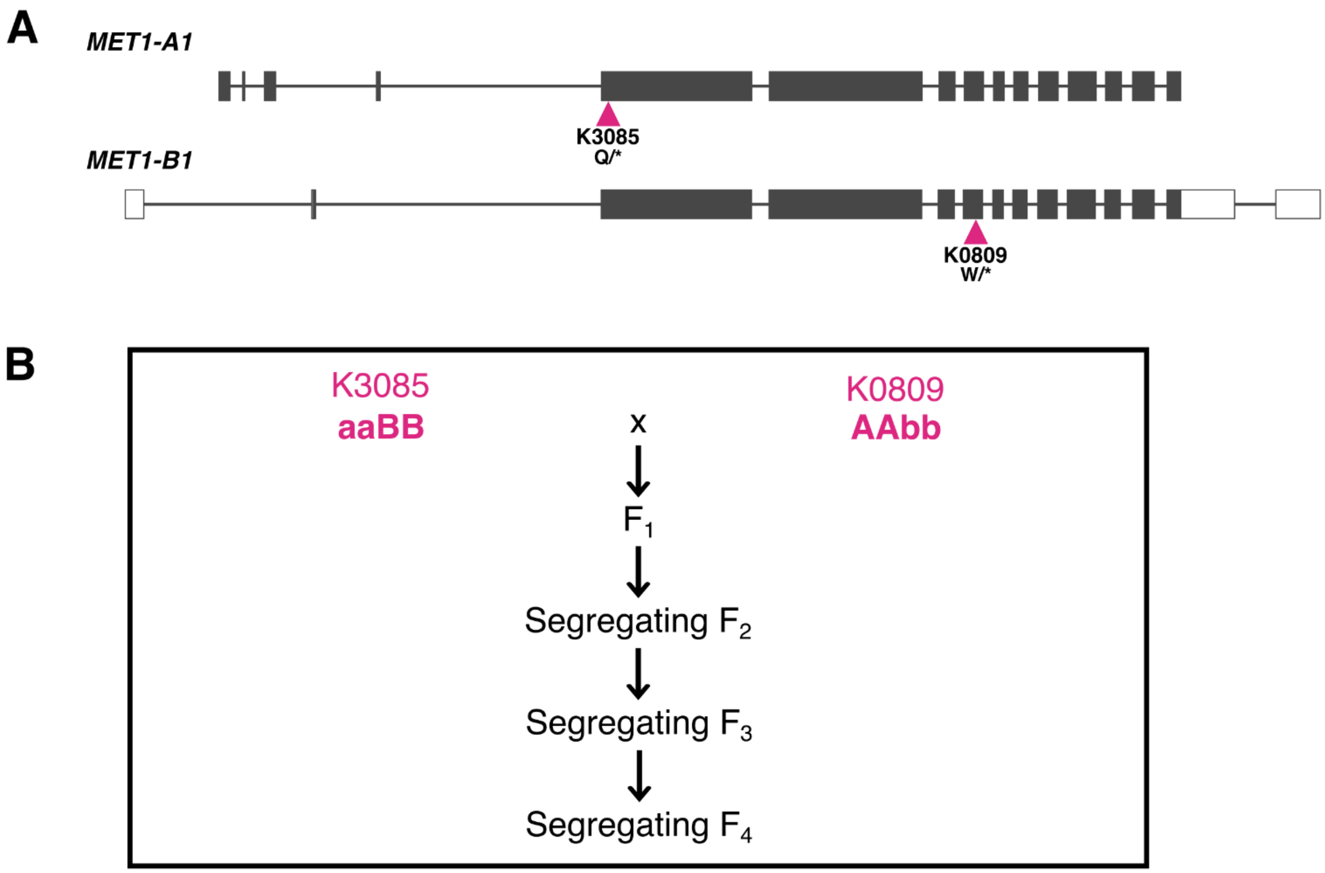


**Supplementary Figure 3.** Overview of the tetraploid *met1-1* TILLING mutants and how they were used to generate mutant populations. A) Structure of the two *MET1-1* homoeologues. Filled rectangles represent exons, empty rectangles represent untranslated regions, and lines represent introns. Triangles indicate the positions of the PTC mutations in the TILLING lines used to produce the segregating populations. B) Crossing structure showing the formation of the segregating populations in the $F_2$, $F_3$ and $F_4$ generations.

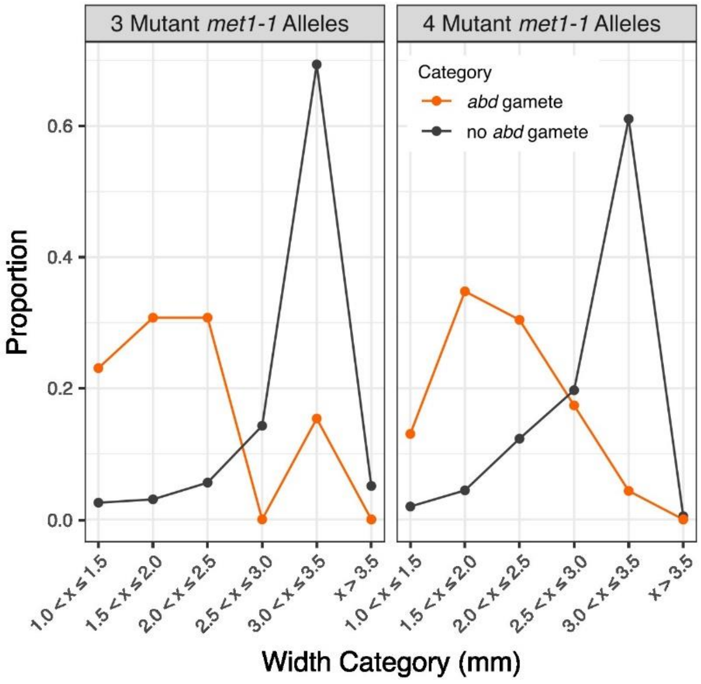


**Supplementary Figure 4.** The proportion of grains in each width category, for grains with a total of three or four mutant *met1-1* alleles, formed from an abd gamete (orange) or not (grey).

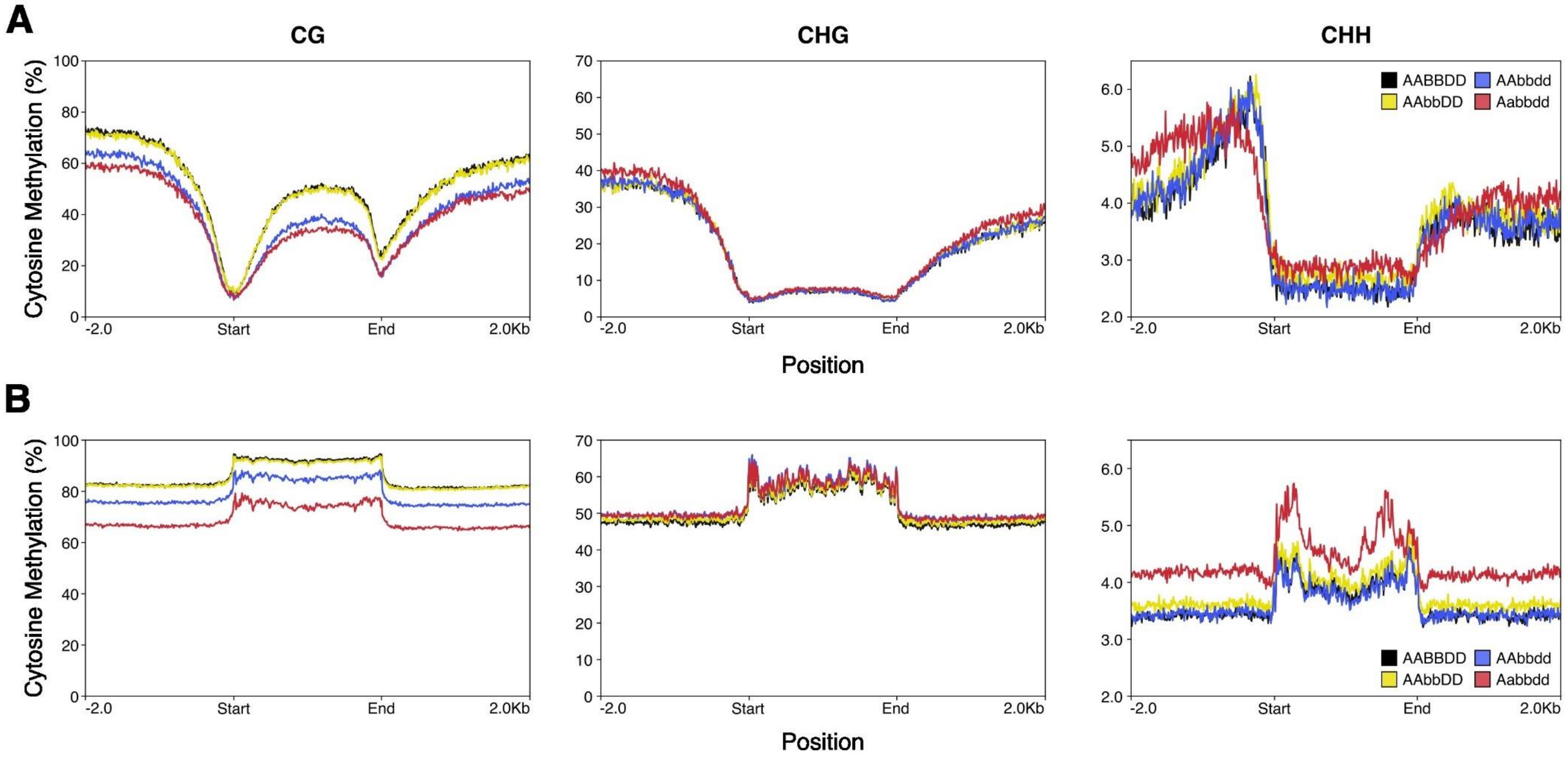


**Supplementary Figure 5.** Percentage cytosine methylation across genes (A) and transposable elements (B) on chromosome 5D, including 2 kb up and downstream of the feature. Cytosine methylation in the CG (left), CHG (centre), and CHH (right) contexts are shown. The WT segregant (AABBDD, black), AAbbDD single mutant (yellow), AAbbdd double mutant (blue), and Aabbdd mutant (red) are shown.

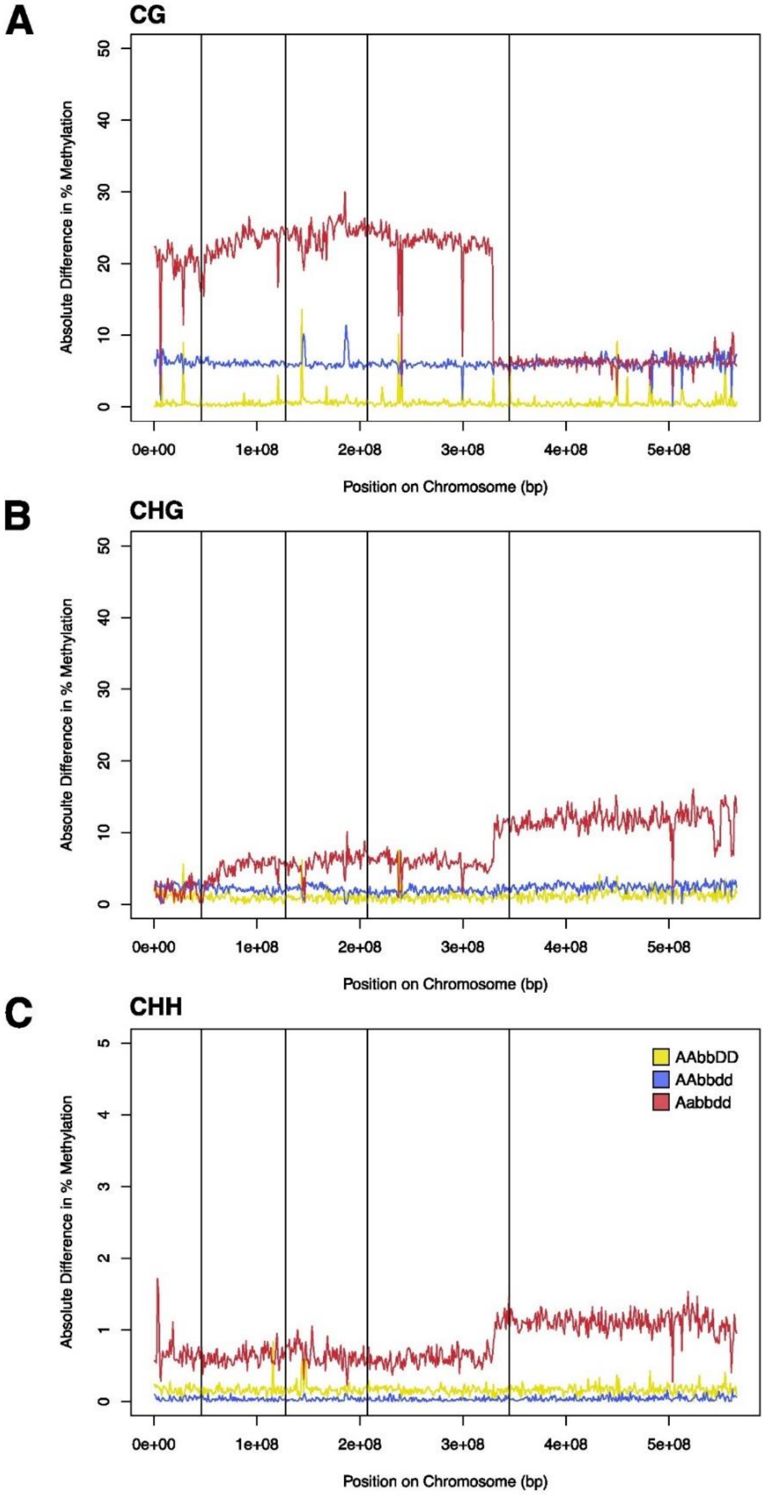


**Supplementary Figure 6.** Absolute difference in percentage methylation between the WT segregant (AABBDD) and the AAbbDD mutant (yellow), the AAbbdd mutant (blue) and the Aabbdd mutant (red) across Chromosome 5D, calculated across 1 Mb bins. Methylation in the CG (A), CHG (B), and CHH (C) contexts are shown. Vertical lines represent the boundaries between chromosomal regions R1 (leftmost), R2a, C, R2b and R3 (rightmost).

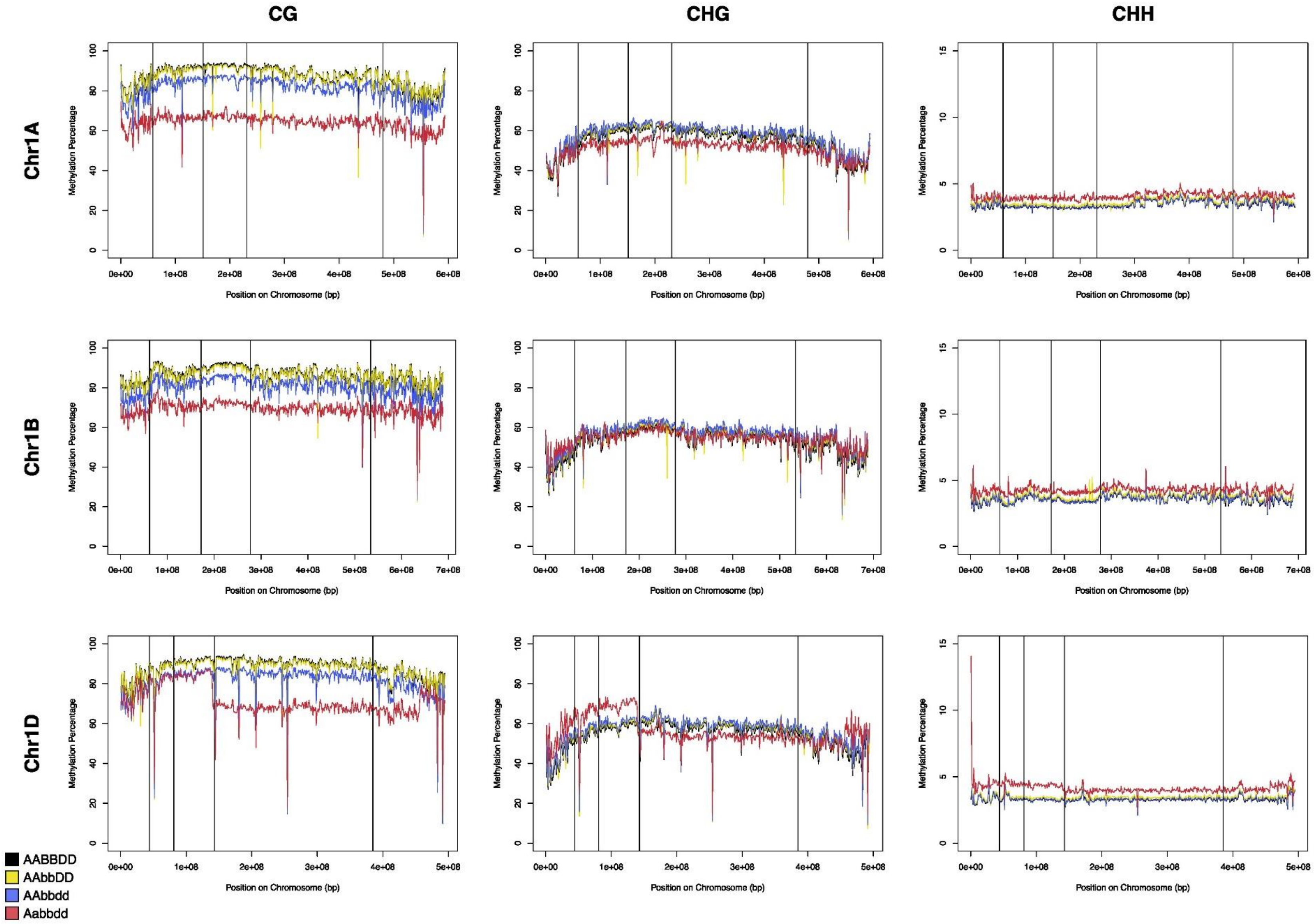


**Supplementary Figure 7.** Percentage methylated cytosines (in the CG, CHG and CHH contexts) calculated for 1 Mb bins across the group 1 chromosomes for the AABBDD genotype (black) and the AAbbDD (yellow), AAbbdd (blue) and Aabbdd (red) mutants. Vertical lines represent the boundaries between chromosomal regions R1 (leftmost), R2a, C, R2b and R3 (rightmost).

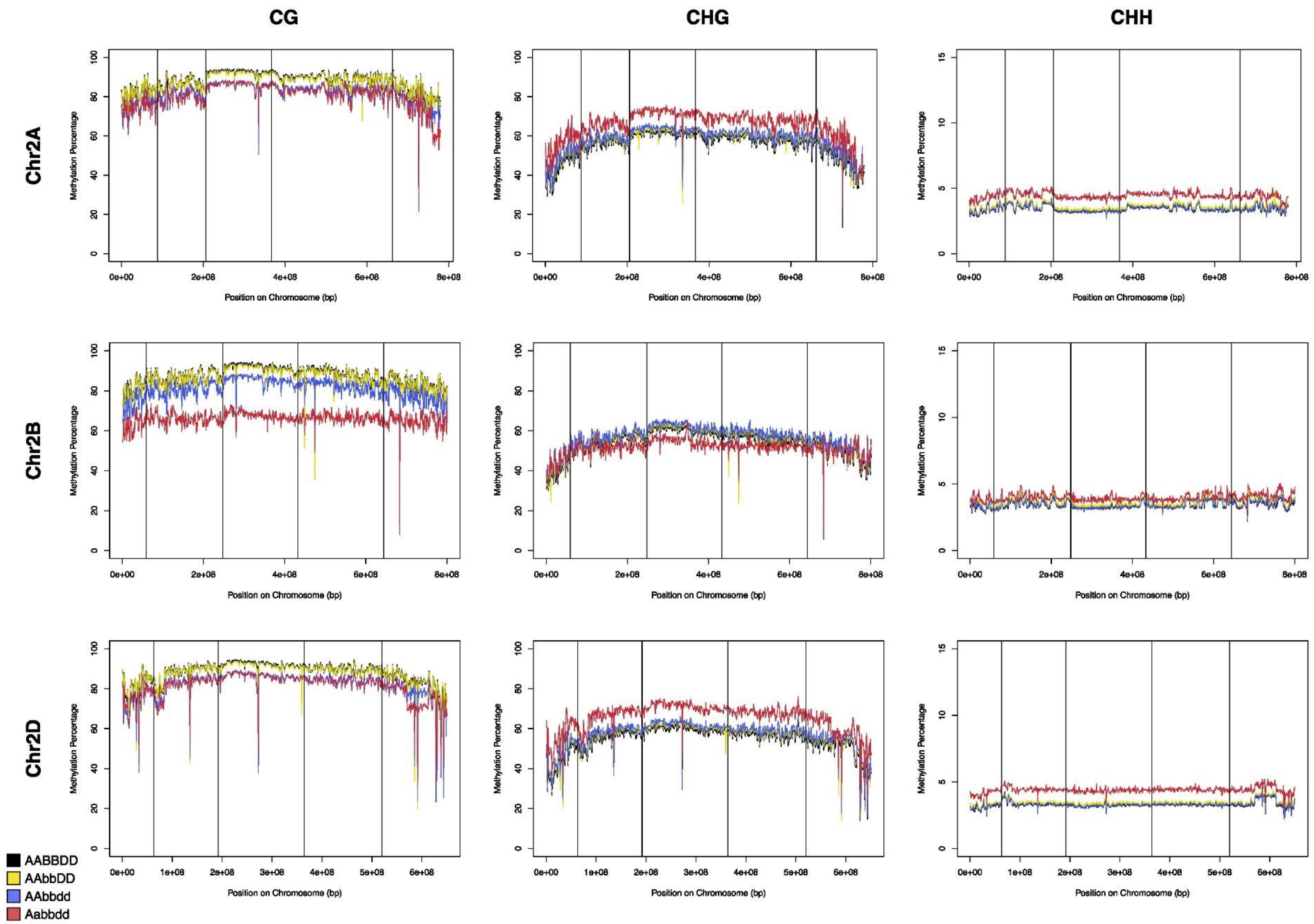


**Supplementary Figure 8.** Percentage methylated cytosines (in the CG, CHG and CHH contexts) calculated for 1 Mb bins across the group 2 chromosomes for the AABBDD genotype (black) and the AAbbDD (yellow), AAbbdd (blue) and Aabbdd (red) mutants. Vertical lines represent the boundaries between chromosomal regions R1 (leftmost), R2a, C, R2b and R3 (rightmost).

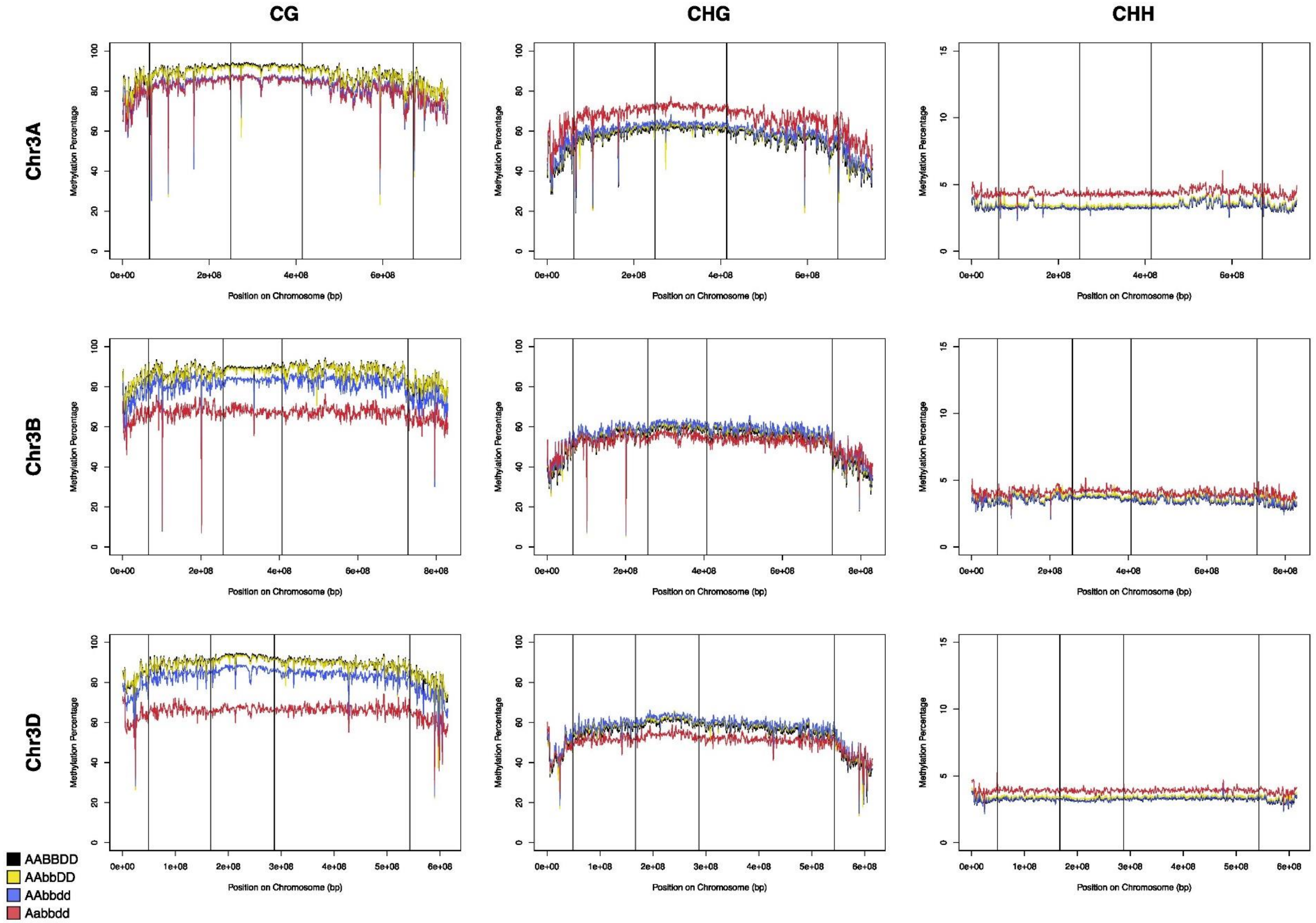


**Supplementary Figure 9.** Percentage methylated cytosines (in the CG, CHG and CHH contexts) calculated for 1 Mb bins across the group 3 chromosomes for the AABBDD genotype (black) and the AAbbDD (yellow), AAbbdd (blue) and Aabbdd (red) mutants. Vertical lines represent the boundaries between chromosomal regions R1 (leftmost), R2a, C, R2b and R3 (rightmost).

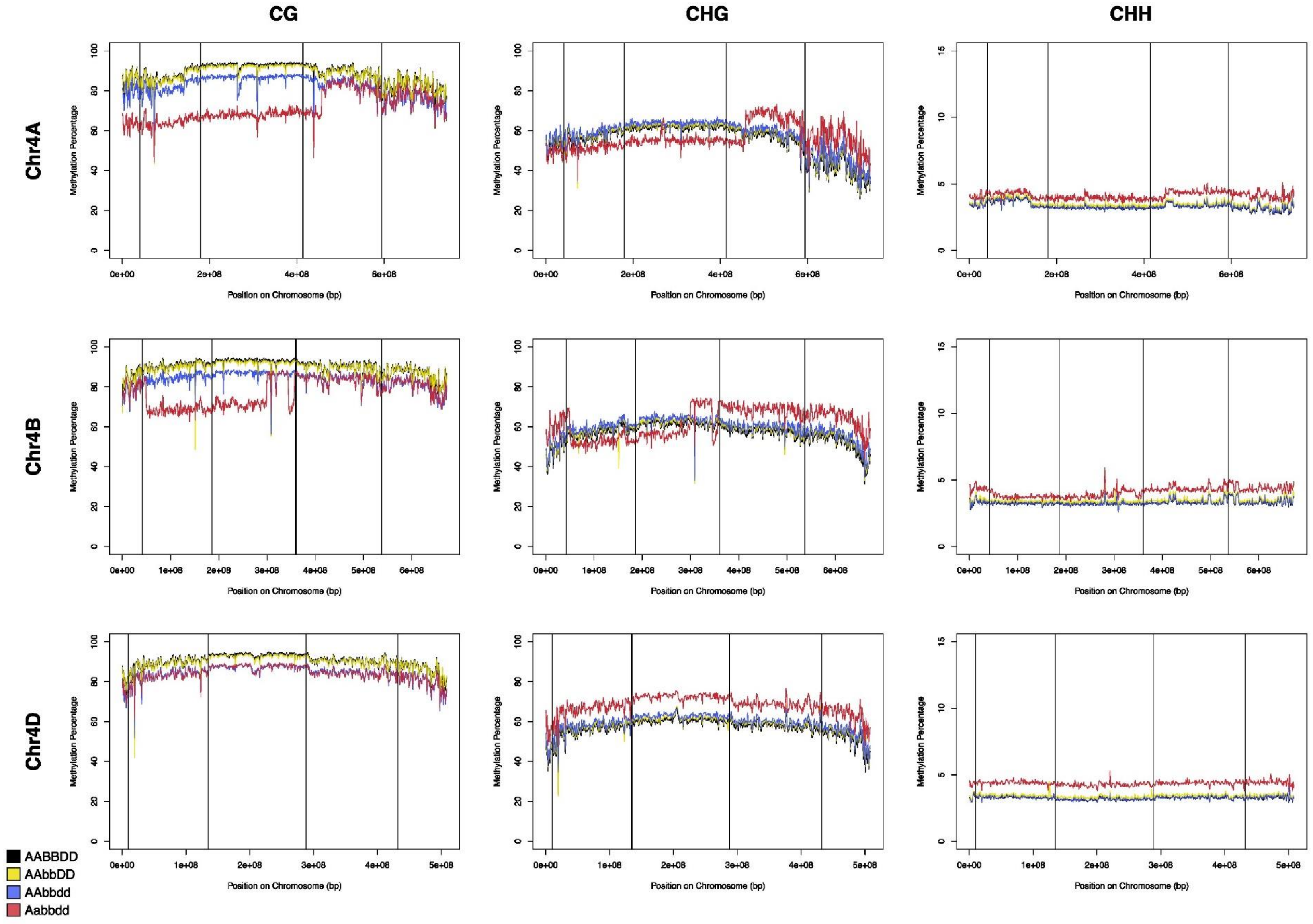


**Supplementary Figure 10.** Percentage methylated cytosines (in the CG, CHG and CHH contexts) calculate for 1 Mb bins across the group 4 chromosomes for the AABBDD genotype (black) and the AAbbDD (yellow), AAbbdd (blue) and Aabbdd (red) mutants. Vertical lines represent the boundaries between chromosomal regions R1 (leftmost), R2a, C, R2b and R3 (rightmost).

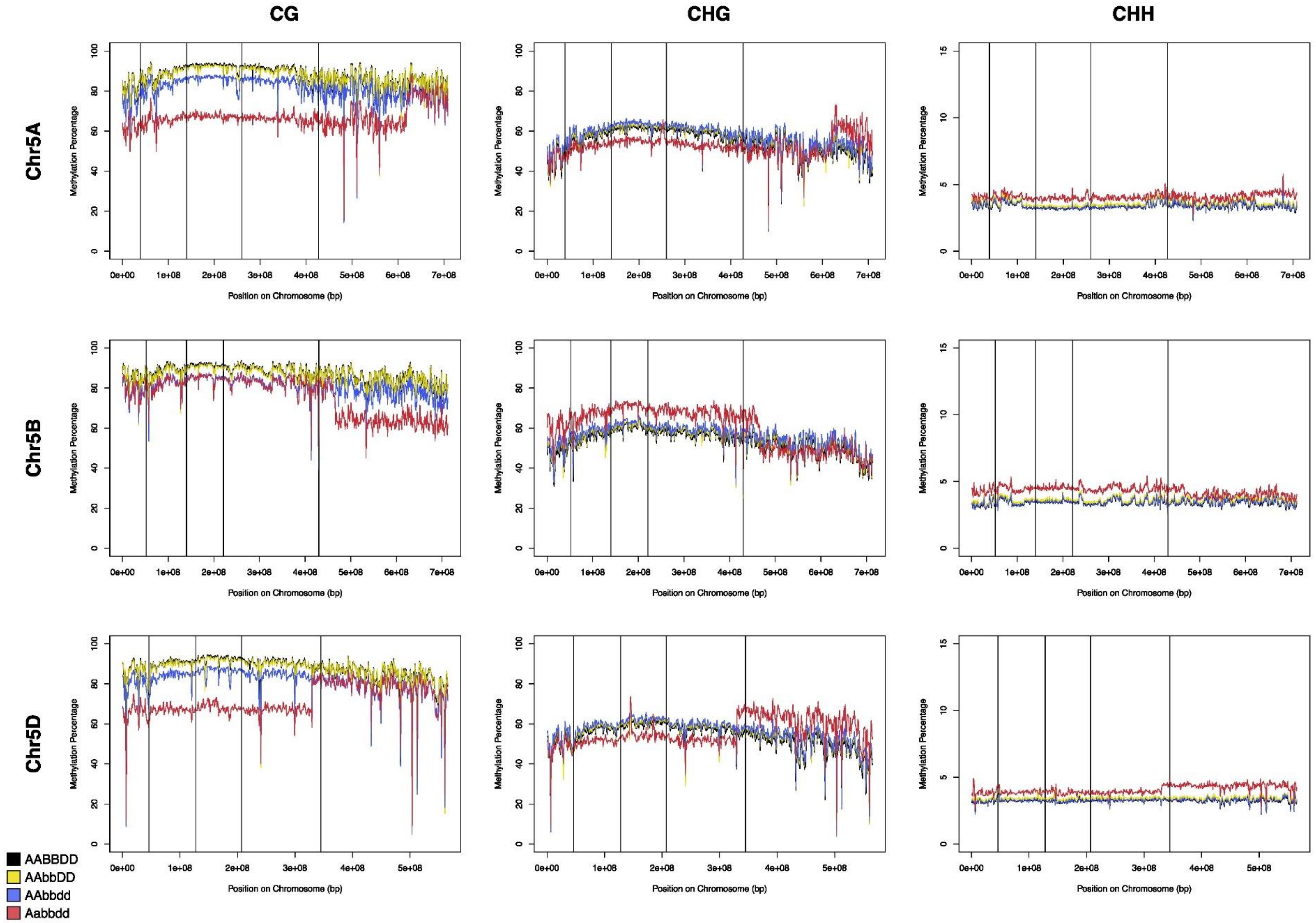


**Supplementary Figure 11.** Percentage methylated cytosines (in the CG, CHG and CHH contexts) calculated for 1 Mb bins across the group 5 chromosomes for the AABBDD genotype (black) and the AAbbDD (yellow), AAbbdd (blue) and Aabbdd (red) mutants. Vertical lines represent the boundaries between chromosomal regions R1 (leftmost), R2a, C, R2b and R3 (rightmost).

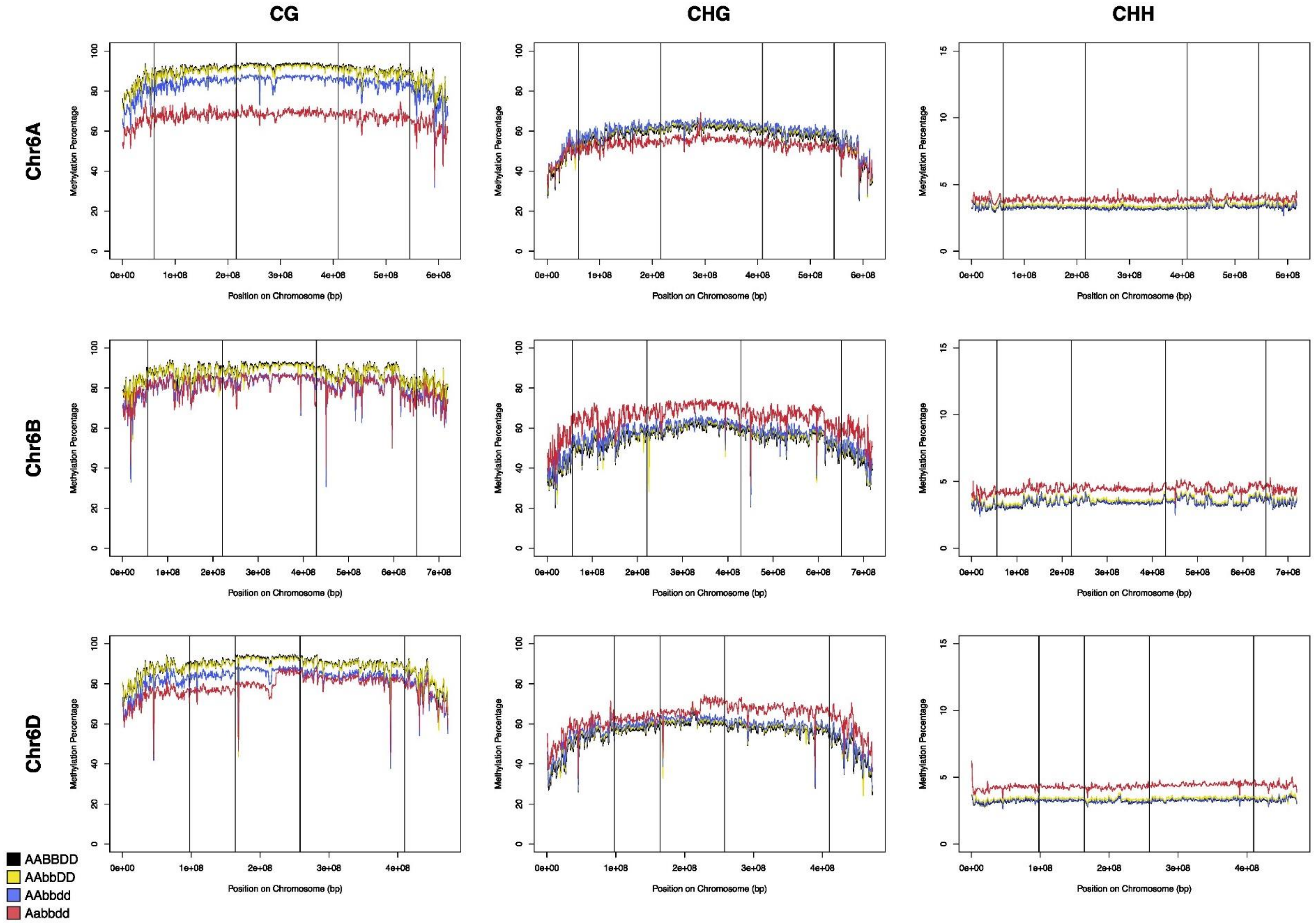


**Supplementary Figure 12.** Percentage methylated cytosines (in the CG, CHG and CHH contexts) calculated for 1 Mb bins across the group 6 chromosomes for the AABBDD genotype (black) and the AAbbDD (yellow), AAbbdd (blue) and Aabbdd (red) mutants. Vertical lines represent the boundaries between chromosomal regions R1 (leftmost), R2a, C, R2b and R3 (rightmost).

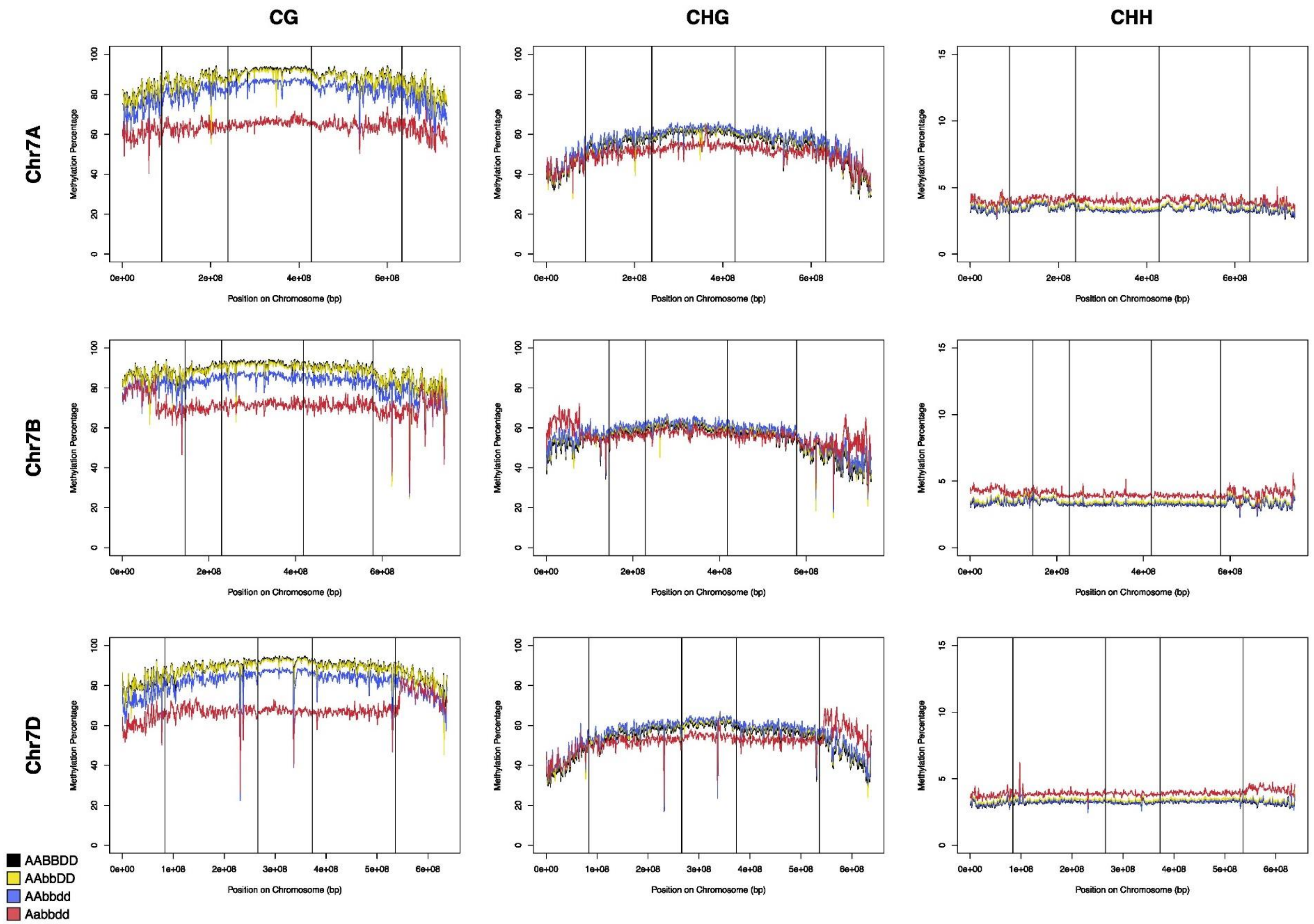


**Supplementary Figure 13.** Percentage methylated cytosines (in the CG, CHG and CHH contexts) calculated for 1 Mb bins across the group 7 chromosomes for the AABBDD genotype (black) and the AAbbDD (yellow), AAbbdd (blue) and Aabbdd (red) mutants. Vertical lines represent the boundaries between chromosomal regions R1 (leftmost), R2a, C, R2b and R3 (rightmost).

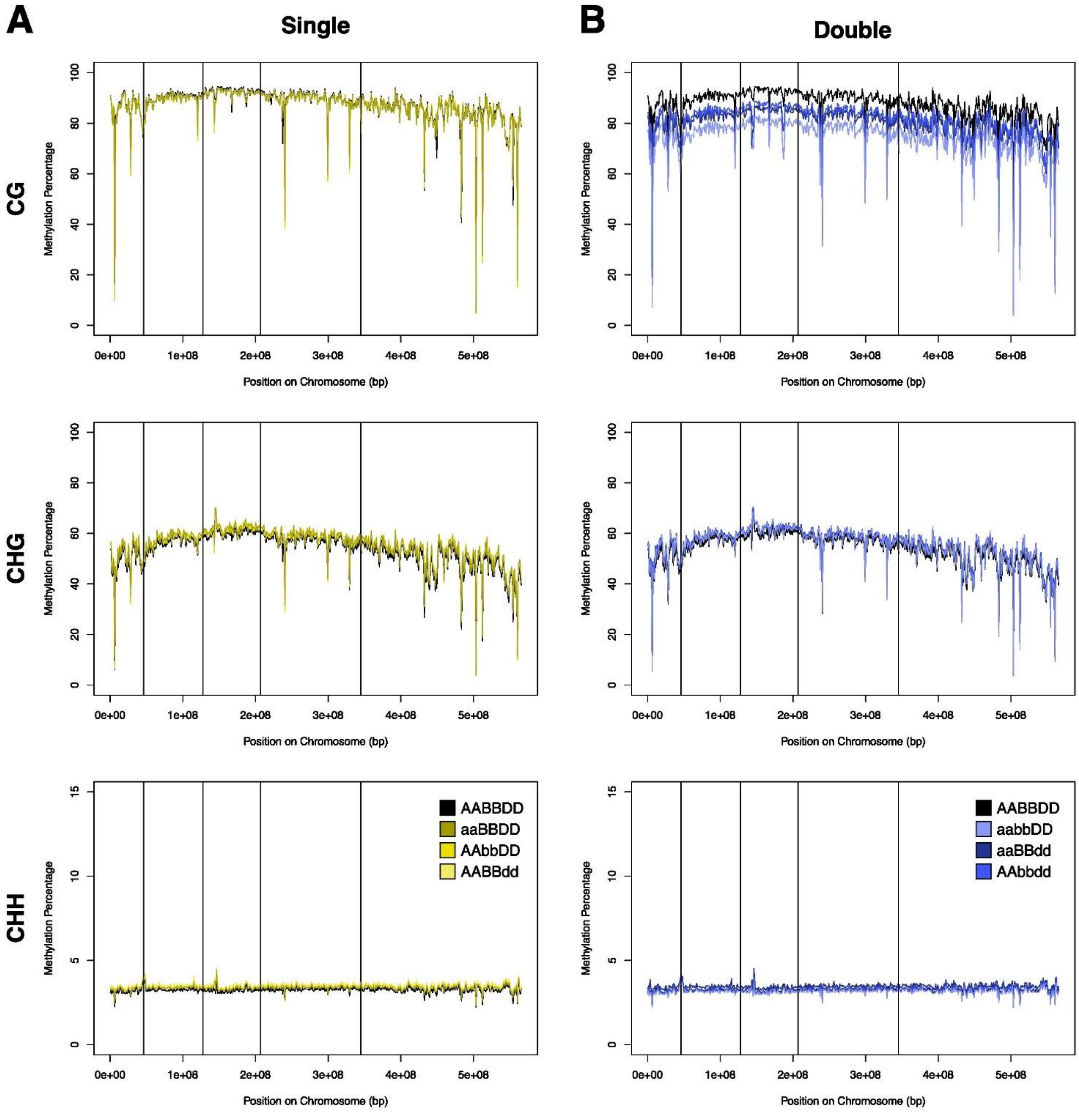


**Supplementary Figure 14.** Percentage of methylated cytosines in the CG, CHG, and CHH contexts across Chromosome 5D for each of the single mutants (A) and double mutants (B), calculated as an average in 1 Mb bins across the chromosome. Vertical lines represent the boundaries between chromosomal regions R1 (leftmost), R2a, C, R2b and R3 (rightmost).

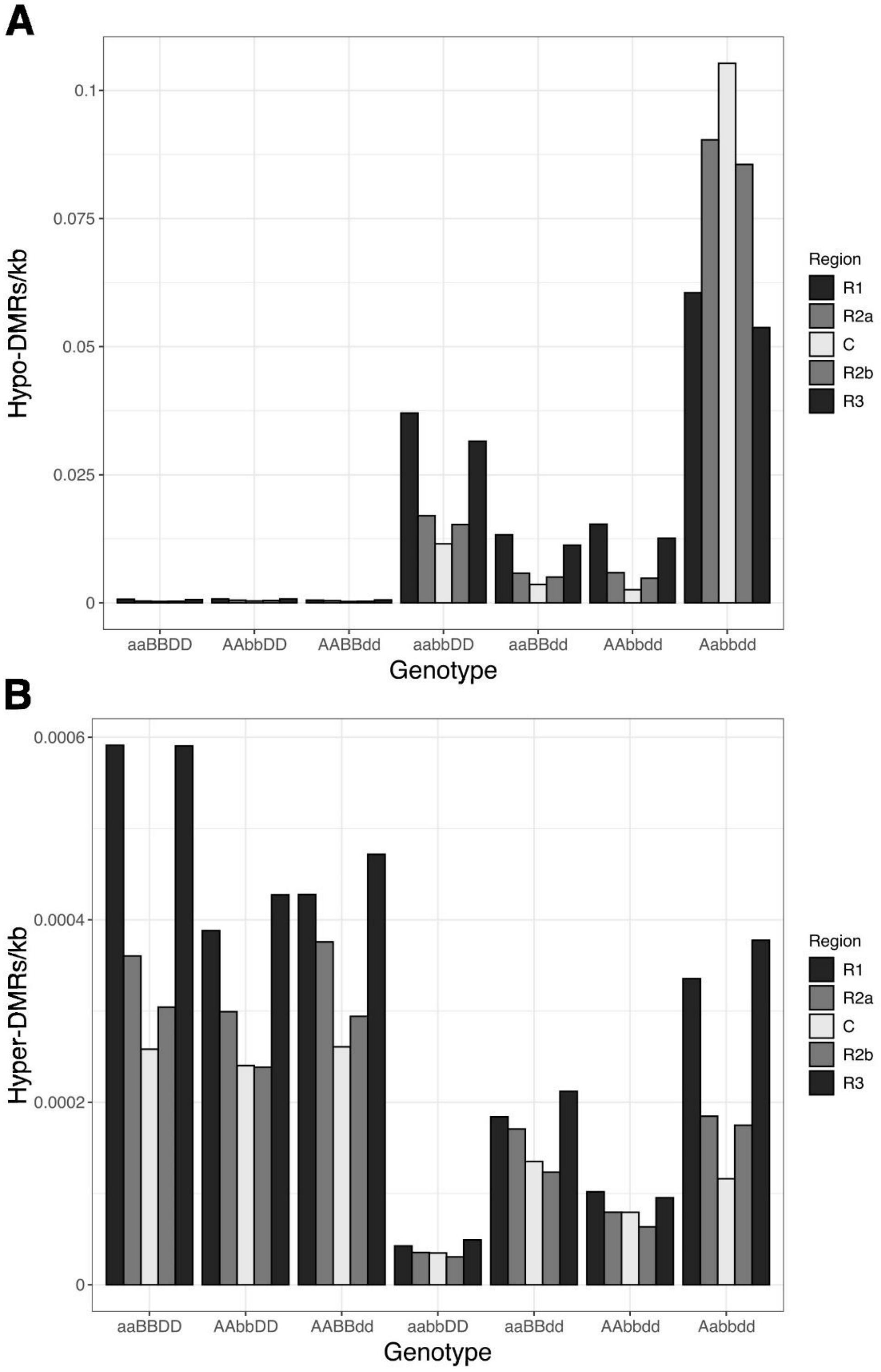


**Supplementary Figure 15.** Number of hypo-methylated (A) and hyper-methylated (B) DMRs per kb in each of the chromosome regions – the distal regions R1 and R3 are shown in dark grey, the proximal regions R2a and R2b in medium grey, and the centromeric region C in light grey.

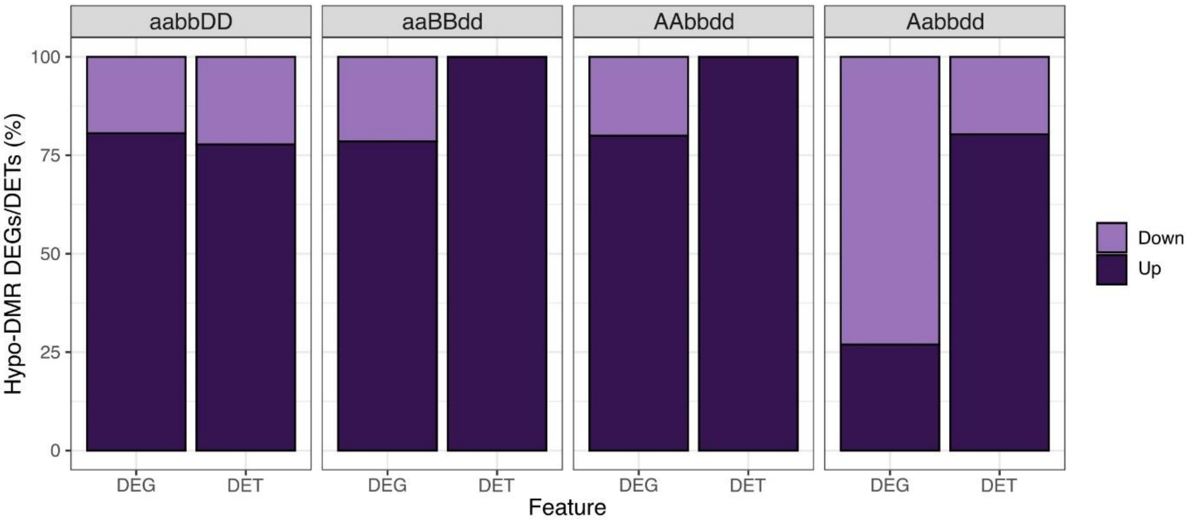


**Supplementary Figure 16.** The percentage of differentially expressed genes (DEGs) and differentially expressed transposons (DETs) associated with a hypo-DMR (differentially methylated region) which are down-regulated (light purple) or up-regulated (dark purple) in the double and Aabbdd *met1-1* mutants.

#### Supplementary Tables

**Supplementary Table 1.** Segregation distortion of *met1-1* genotypes by mutant copy number in the C0451xC2028xC1292 population $F_2$ generation.

| Mutant copy No. | Observed Count | Expected Count |
|---|---|---|
| 0 | 15 | 5.13 |
| 1 | 34 | 30.75 |
| 2 | 98 | 76.88 |
| 3 | 116 | 102.50 |
| 4 | 65 | 76.88 |
| 5 | 0 | 30.75 |
| 6 | 0 | 5.13 |
| **Total** | 328 | 328 |
| **Chi Squared p-value** | $5.05 \times 10^{-12}$ | |

**Supplementary Table 2.** Grain genotyping results validated by leaf genotyping for plants with 3 marker results. Perfect match = genotyping results of all three homoeologs using grain DNA match the results from leaf DNA.

| | Number of plants | Percent match |
|---|---|---|
| **A Match** | 162 | 100.0% |
| **B Match** | 159 | 98.1% |
| **D Match** | 160 | 98.8% |
| **Perfect Match** | 157 | 96.9% |
| **Total Leaves** | 162 | |

**Supplementary Table 3.** Segregation distortion of Kronos $F_4$ grain genotypes by mutant copy number in selfed double heterozygous (AaBb) plant offspring. Mutant *met1-1* alleles were classified as paternally or maternally inherited by grain genotyping.

| Mutant copy No. | Observed Count | Expected Count |
|---|---|---|
| 0 | 44 | 20.75 |
| 1 | 135 | 83.00 |
| 2 | 142 | 124.50 |
| 3 | 11 | 83.00 |
| 4 | 0 | 20.75 |
| **Total** | 332 | 332 |
| **Chi Squared p-value** | $3.39 \times 10^{-30}$ | |

| No. Paternal Mutant Copies | Observed Count | Expected Count |
|---|---|---|
| 0 | 101 | 83 |
| 1 | 213 | 166 |
| 2 | 18 | 83 |
| **Total** | 332 | 332 |

| No. Maternal Mutant Copies | Observed Count | Expected Count |
|---|---|---|
| 0 | 130 | 83 |
| 1 | 201 | 166 |
| 2 | 1 | 83 |
| **Total** | 332 | 332 |

**Supplementary Table 4**. Pollen phenotypes are not significantly different between *met1-1* mutants and AABBDD plants. The estimated marginal means for each phenotype are shown for each genotype category. Shared letters show genotypes that are not significantly different from one another (FDR adjusted p<0.05).

| Genotype Category | Pollen Grains per Anther | Modal Pollen Diameter (μm) | Non-Viable Pollen Grains (%) | Non- and Mononucleated Pollen Grains (%) |
|---|---|---|---|---|
| Cadenza WT | 2,459 (a) | 49.8 (a) | 1.05 (a) | 8.41 (a) |
| AABBDD | 1,829 (bc) | 46.2 (b) | 2.62 (b) | 12.45 (ab) |
| aaBBDD/AAbbDD/AABBdd | 2,085 (ab) | 46.8 (b) | 1.96 (b) | - |
| aabbDD/aaBBdd/AAbbdd | 1,623 (c) | 45.3 (b) | 1.60 (ab) | 11.40 (a) |
| AaBbdd/AabbDd/aaBbDd | 1,780 (bc) | 45.8 (b) | 2.10 (b) | 19.56 (b) |
| N per genotype category | 10-15 | 10-16 | 16-38 | 4-15 |

**Supplementary Table 5.** Transposable elements (TEs) that intersect at least one differentially methylated region (DMR), categorised according to class and family, as a percentage of the total number of TEs that overlap a DMR in each mutant genotype. Class I TEs include long terminal repeat (LTR) retrotransposons which consist largely of Copia and Gypsy elements. Class II TEs consist largely of terminal inverted repeat (TIR) transposons. TEs which are not Class I or Class II are unclassified.

| | | | **aaBBDD** | **AAbbDD** | **AABBdd** | **aabbDD** | **aaBBdd** | **AAbbdd** | **Aabbdd** |
|---|---|---|---|---|---|---|---|---|---|
| **Class I** | | | **81.0** | **82.0** | **84.8** | **74.7** | **74.2** | **60.0** | **88.2** |
| | LTR | | 79.3 | 80.2 | 83.2 | 72.5 | 72.2 | 56.5 | 87.0 |
| | | Copia | 16.7 | 15.6 | 16.9 | 18.4 | 17.3 | 12.5 | 20.5 |
| | | Gypsy | 54.7 | 57.0 | 59.6 | 47.8 | 48.3 | 36.5 | 61.3 |
| **Class II** | | | **13.5** | **13.6** | **11.8** | **18.5** | **18.7** | **28.9** | **9.5** |
| | TIR | | 13.5 | 13.6 | 11.5 | 18.4 | 18.6 | 28.6 | 9.3 |
| **Unclassified** | | | **5.5** | **4.4** | **3.4** | **6.8** | **7.0** | **11.1** | **2.3** |

**Supplementary Table 6.** Primer sequences used for KASP genotyping and PCR.

| **Name** | **Sequence (5'-3')** | **Purpose** |
|---|---|---|
| Met1_2A_Cad0465_WT | GAAGGTCGGAGTCAACGGATTaatgaagttgattcaggcaaatG | KASP genotyping Cadenza C0465 x C0884 *MET1-1* population |
| Met1_2A_Cad0465_Mut | GAAGGTGACCAAGTTCATGCTaatgaagttgattcaggcaaatA | |
| Met1_2A_Cad0465_Com | ACAGCCAGCAAAAATGTCG | |
| Met1_2B_Cad0465_WT | GAAGGTCGGAGTCAACGGATTcccatacagacccttccactG | |
| Met1_2B_Cad0465_Mut | GAAGGTGACCAAGTTCATGCTcccatacagacccttccactA | |
| Met1_2B_Cad0465_Com | GTGTTTGTGAGATGTTAATTCCCA | |
| Met1_2D_Cad0884_WT1 | GAAGGTCGGAGTCAACGGATTaaatatattggaactcctacatcctG | |
| Met1_2D_Cad0884_Mut1 | GAAGGTGACCAAGTTCATGCTaaatatattggaactcctacatcctA | |
| Met1_2D_Cad0884_Com1 | TGTCTTTGGATCTGGTTTTATGAC | |
| Met1_2A_Cad0451_WT | GAAGGTCGGAGTCAACGGATTcccctctgtttcttcatcatcttC | KASP genotyping Cadenza C0451 x C2028 x C1292 *MET1-1* population |
| Met1_2A_Cad0451_Mut | GAAGGTGACCAAGTTCATGCTcccctctgtttcttcatcatcttT | |
| Met1_2A_Cad0451_Com | AGTGTAAAGAGGTAGCTGAGGAT | |
| Met1_2D_Cad2028_WT | GAAGGTCGGAGTCAACGGATTttgggcgtattgaggactgG | |
| Met1_2D_Cad2028_Mut | GAAGGTGACCAAGTTCATGCTttgggcgtattgaggactgA | |
| Met1_2D_Cad2028_Com | GTTCTGTTGAGAGCCAGACT | |
| Met1_2B_Cad1292_F | GCAACAAATGGTGTTGAAAAGT | CAPS genotyping Cadenza C0451 x C2028 x C1292 *MET1-1* population |
| Met1_2B_Cad1292_R | GCCTTCTCATAAAAGTGGTTGA | |
| MET1-A_K3085_WT_H | GAAGGTCGGAGTCAACGGATTaggtgcagagcaaagccaC | KASP genotyping Kronos *MET1-1* population |
| MET1-A_K3085_M_F | GAAGGTGACCAAGTTCATGCTaggtgcagagcaaagccaT | |
| MET1-A_K3085_C | ccaattttccttgtgcggtgA | |
| MET1-B_K0809_WT_H | GAAGGTCGGAGTCAACGGATTctctctttgggcacTtccatC | |
| MET1-B_K0809_M_F | GAAGGTGACCAAGTTCATGCTctctctttgggcacTtccatT | |
| MET1-B_K0809_C | tcatgtctgtggctcgcaag | |

**Supplementary Methods**

*Pollen Assays*

Plants used for the pollen assays were grown in a randomised block design. Number of pollen grains per anther and pollen grain diameter were analysed using the Multisizer 4e Coulter Counter (Beckman Coulter, California, USA). Per plant, three non-extruded, mature anthers were collected from three florets on the secondary spike. Anthers were stored in 70% ethanol at 4°C. Pollen grains were released and analysed as previously described (Alabdullah *et al.*, 2021). Two measurements were performed per floret sample; one measuring all pollen grains in a total volume of 2 mL (to calculate mean pollen grains per anther), and another measuring up to 10,000 particles (to calculate modal pollen diameter). To remove the effect of debris, only particles with diameter between 30 and 60 µm were considered for this analysis.

Pollen viability was analysed using the Ampha Z32 Pollen Analyzer (Amphasys, Lucerne, Switzerland). 10 non-extruded, mature anthers were collected per plant from the primary spike and ruptured to release the pollen grains into AmphaFluid9 buffer (Amphasys). Up to 10,000 pollen grains were measured per sample; samples with <2,000 pollen grains were excluded from the analysis. To identify the cluster corresponding to non-viable pollen grains, anthers from Cadenza WT plants were heated to 65°C to kill the pollen. Separately, we agitated a sample of AmphaFluid9 buffer to identify the cluster corresponding to bubbles. Linear mixed models were used to calculate the estimated marginal means for pollen number per anther, pollen size, and percentage pollen viability, per genotype category, with block as a random variable (phenotype ~ genotype category + (1|block)). Pairwise significant differences were assigned using a false discovery rate multiple testing correction. Statistical analysis was carried out using the lme4 and emmeans packages in R (v4.2.2).

To determine the number of nuclei per pollen grain, pollen grains were stained with DAPI (4',6-diamidine-2-phenylindole). Four non-extruded, mature anthers from the primary spike were fixed in Carnoy's solution. After a minimum incubation of 48 hours, the anthers were removed from the Carnoy's solution and stained with DAPI solution (0.001 mg/mL DAPI, 1% (v/v) Triton X-100). Pollen grains were released by sonication and incubated in the dark for 5 minutes. Samples were imaged with an Axio Zoom v16 II stereomicroscope (Zeiss, Oberkochen, Germany) with ORCA-Flash4.0 Digital CMOS camera (Hamamatsu Photonics, Shizuoko, Japan). Six non-overlapping images were collected per sample. Non-, mono-, bi-, and tri-nucleated pollen grains were counted for each sample. Pollen grains with ambiguous numbers of nuclei were disregarded. As fewer replicates were used for DAPI staining, estimated marginal means were calculated irrespective of blocks and significant differences assigned by a Wilcoxon signed-rank test with false discovery rate multiple testing correction. The percentage of pollen grains with abnormal numbers of nuclei were log transformed before the test was applied.
